# DHX36 regulates antral follicle development and ovulation as a non-OSF maternal factor by maintaining oocyte homeostasis and supporting OSF delivery

**DOI:** 10.64898/2026.08.18.745561

**Authors:** Yu-Xuan Jiao, Fang-Yin Sun, Guo-Wei Bu, Yu-Ling Chen, Kunpeng Zhou, Bo-Ya Guo, Hai-Teng Deng, Yi-Zhen Sima, Hong-Ying Sha, Su-Ying Liu, Yong-Juan Sang, Qi-Ming Sun, Xiaona Chen, Huating Wang, Cunqi Ye, Heng-Yu Fan

**Author notes:** Correspondence: Heng-Yu Fan. Co-first authors (Yu-Xuan Jiao, Fang-Yin Sun, Guo-Wei Bu, Yu-Ling Chen).

## Abstract

Healthy ovarian follicle development and ovulation require coordinated communication between oocytes and surrounding somatic cells. Although oocyte-secreted factors (OSFs), such as GDF-9 and BMP-15, are established regulators of this communication, the non-OSF maternal factors that support OSF delivery and signaling during late-stage follicle development remain poorly understood. Here, using an oocyte-specific *Dhx36* knockout mouse model, we identify the G-quadruplex (G4) helicase DHX36 as a non-OSF maternal factor required for antral follicle development and hormone-induced ovulation. *Dhx36* deficiency caused severe defects in granulosa cell proliferation and cumulus expansion, accompanied by impaired activation of SMAD2/3 and SMAD1/5/8, while ERK1/2 activation remained intact. Although the expression of major OSFs was largely unchanged, *Dhx36*-deficient oocytes exhibited disrupted microvilli and transzonal projections (TZPs), resulting in defective OSF delivery and impaired oocyte–cumulus communication. Proteomic, lipidomic, and ultrastructural analyses further revealed dysregulated phospholipid metabolism, membrane organization, autophagy, and organelle homeostasis, including abnormal lysosomal, mitochondrial, and endoplasmic reticulum structures. Integrative transcriptomic and proteomic analyses identified concordant downregulation of genes involved in these processes, whose promoters were enriched in potential G4 motifs. Consistently, *Dhx36* deficiency was associated with reduced RNA polymerase II activity, while pharmacological G4 stabilization impaired transcription of selected genes. Together, these findings establish DHX36 as a maternal regulator that links oocyte intrinsic homeostasis to intercellular communication, suggesting that DHX36-dependent maintenance of membrane and organelle integrity is essential for OSF delivery, cumulus cell function, antral follicle development, and ovulation.

## Introduction

Healthy ovarian follicle development and successful ovulation are prerequisites for female fertility^1^. Following activation of primordial follicles, follicles progressively develop through the primary and secondary stages to form antral follicles, which subsequently undergo ovulation and release the cumulus–oocyte complex (COC) into the oviduct^2–6^.

Oocyte-secreted factors (OSFs), particularly growth differentiation factor 9 (GDF-9) and bone morphogenetic protein 15 (BMP-15), are well-established regulators of follicle development and ovulation^7^. For example, maternal deletion of *Gdf9*, causes follicular arrest at primary stage^8^. Efficient OSF delivery from oocytes to surrounding cumulus cells requires specialized structures, including oocyte microvilli and transzonal projections (TZPs) from granulosa cells^9^. Impaired microvilli or TZPs can reduce GDF-9 delivery and compromise follicle development and female fertility^10^. In cumulus cells, OSFs activate TGFβ/SMAD signaling to induce downstream genes, including *Cox2*, *Has2*, *Tnfaip6*, and *Ptx3*, thereby promoting cumulus expansion and ovulation^11^. M Meanwhile, LH-induced ErbB–ERK1/2 signaling also contributes to these processes^12^.

Increasing evidence indicates that non-OSF maternal factors, including *Rtcb*^13^, *Pabpn1*^14^, and *Dcaf13*^15^, are also indispensable for follicle development, as their deletion causes a marked reduction in follicle number from the primary to secondary stages and ultimately leads to severe primary ovarian insufficiency (POI). However, few non-OSF maternal factors have been identified that regulate later-stage follicle development, particularly antral follicle development and ovulation, or whose functions are linked to OSF delivery and signaling.

DHX36 is a G-quadruplex (G4) helicase that regulates gene expression^16,17^. Previous study using an oocyte-specific *Zp3*-Cre conditional knockout mouse model showed that *Dhx36* deletion had little effect on ovarian morphology at 8–10 weeks of age but progressively reduced hormone-induced ovulation from 4 weeks of age onward^18^. These findings suggest that maternal DHX36 may have a specific role in later-stage antral follicle development and ovulation.

Here, using integrated histological, proteomic, and lipidomic analyses, we identify DHX36 as a non-OSF maternal factor required for antral follicle development and hormone-induced ovulation. We show that loss of oocyte DHX36 disrupts microvilli structure and oocyte cellular homeostasis, thereby impairing OSF delivery and downstream signaling in cumulus cells and ultimately compromising granulosa cell proliferation, cumulus expansion, and ovulation.

## Results

### Oocyte-specific loss of *Dhx36* suppresses antral follicle development and granulosa cell proliferation in antral follicle

Oocyte-specific *Dhx36* knockout (*Dhx36^oo-/-^*) was generated by crossing *Dhx36^fl/fl^* and *Zp3*-Cre mice, in which Cre-mediated *Dhx36* deletion is initiated in oocyte at the primary follicle stage (**Fig. 1A**). Efficient ablation of DHX36 protein in fully-grown oocytes (FGO) was confirmed by western blotting (**Fig. S1A**). To the role of DHX36 on follicle development and ovulation, we first examined ovaries from 4-month-old *Dhx36^oo-/-^* mice. *Dhx36* deficiency resulted in an age-dependent ovarian insufficiency-like phenotype, characterized by an approximately 30% reduction in ovarian size **(Fig. 1B; Fig. S1B**), an absence of visible corpora lutea (CL) (**Fig. 1C; Fig. S1C**), and a selective reduction in antral follicle without affecting earlier-stage follicles compared with wild-type (WT) littermate controls (**Fig. 1D**). Although antral follicle size was comparable between genotypes (**Fig. S1D**), both cumulus cell number and total granulosa cell number were markedly reduced in *Dhx36^oo-/-^* follicles (**Fig. 1E; Fig. S1E**).

**Figure 1.**
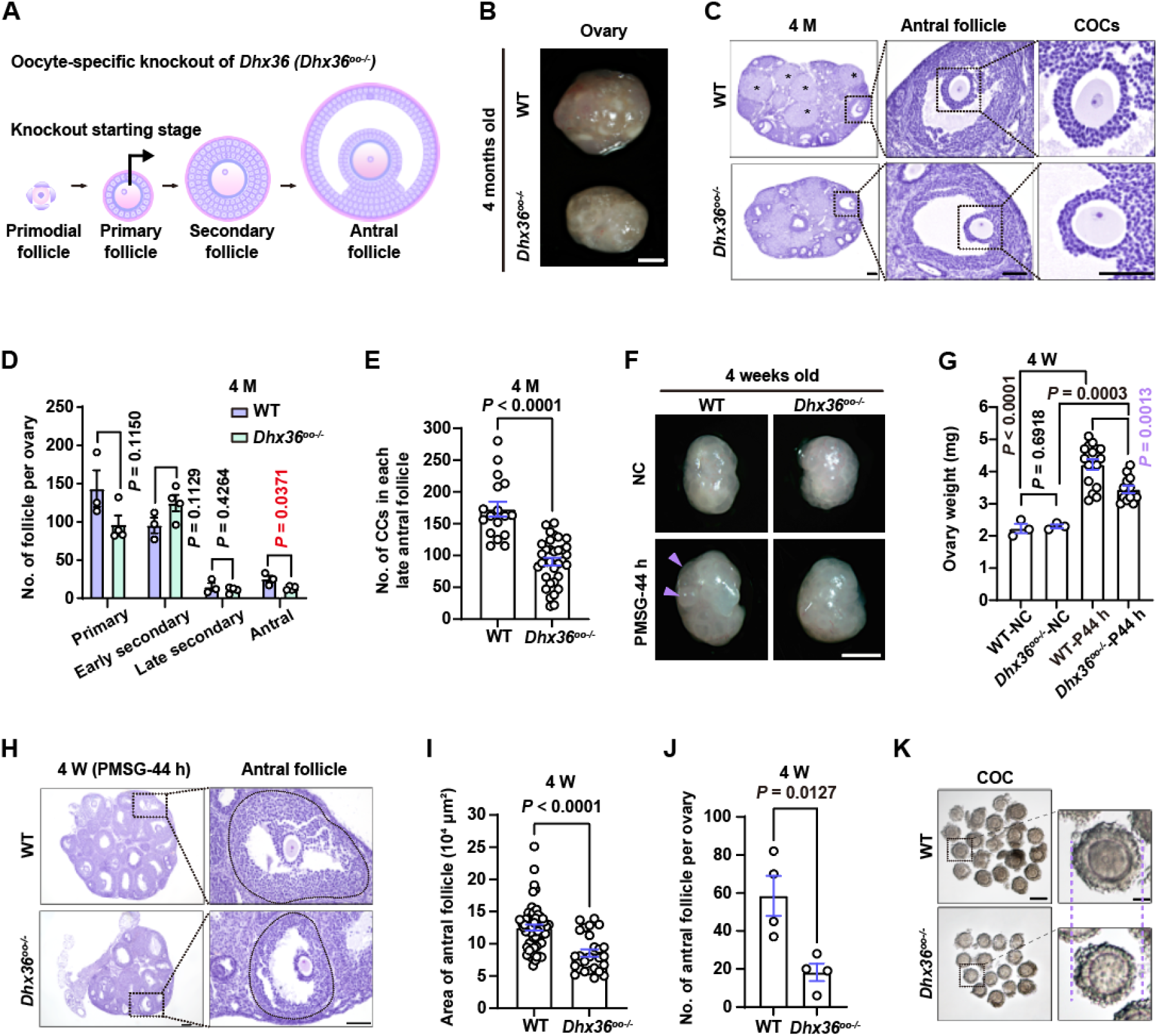
Oocyte-specific deletion of *Dhx36* impairs follicle development A: Schematic diagram illustrating the starting stage of oocyte-specific *Dhx36* knockout. **B:** Representative images of ovaries from 4-month-old WT and *Dhx36^oo-/-^* mice. (Scale bar: 1 mm). **C:** H&E staining images of ovaries from 4-month-old WT and *Dhx36^oo-/-^*mice. Asterisks indicate CL. (Scale bar: 100 μm, 50 μm, and 50 μm, left to right). **D:** Number of follicles at different developmental stages (primary, secondary, and antral) in ovaries from 4-month-old WT and *Dhx36^oo-/-^* mice. **E:** Quantification of CL number per largest ovarian sections from 4-month-old WT and *Dhx36^oo-/-^* mice. α-Tubulin serves as an internal control. **F:** Representative morphology of ovaries from 4-week-old WT and *Dhx36^oo-/-^* mice, either untreated (no treatment control, NC) or treated with PMSG for 44 h. Purple triangles indicate the antral follicles. (Scale bar: 1 mm). **G:** Quantification of ovarian weight in the four experimental groups in (F). **H:** H&E staining images of ovaries from 4-week-old WT and *Dhx36^oo-/-^* mice after PMSG treatment for 44 h. (Scale bar: 100 μm and 50 μm, left to right). **I:** Quantification of antral follicle area in ovaries from WT and *Dhx36^oo-/-^* mice after PMSG treatment for 44 h. **J:** Number of antral follicles in ovaries from WT and *Dhx36^oo-/-^* mice after PMSG treatment for 44 h. **K:** Representative brightfield images of COCs isolated from WT and *Dhx36^oo-/-^* mice after PMSG treatment for 44 h. (Scale bar, left: 100 μm, right: 25 μm). All quantitative data are presented as mean ± SEM. *P* values were calculated using unpaired *t*-tests.

Similar defects were observed in 4-week-old *Dhx36^oo-/-^*mice following gonadotropin stimulation, including reduced ovarian size (**Fig. 1F, G**), decreased antral follicle area (**Fig. 1H, I**), fewer antral follicles per ovary (**Fig. 1J**), and impaired ovulation accompanied by reduced CL formation (**Fig. S1F, G**). Consistently, COCs isolated from *Dhx36^oo-/-^* mice following PMSG stimulation exhibited a markedly thinner cumulus cell layer (**Fig. 1K; Fig. S1H**). Flow cytometric analysis further revealed reductions in both the total granulosa cell number and the proportion of S-phase granulosa cells in antral follicles from *Dhx36^oo-/-^* mice (**Fig. 2A; Fig. S1I, J**). Consistent with these findings, immunofluorescence (IF) and immunohistochemical (IHC) analyses of H3S10ph (**Fig. 2B**–**E**) and IHC analysis of 5-bromo-2’-deoxyuridine (BrdU) incorporation (**Fig. 2F, G**) demonstrated reduced proliferation of both cumulus and mural granulosa cells. These defects were accompanied by reduced serum estradiol (E2) and progesterone (P4) level following PMSG and human chorionic gonadotropin (hCG) treatment, respectively, compared with WT mice (**Fig. 2H, I**).

**Figure 2.**
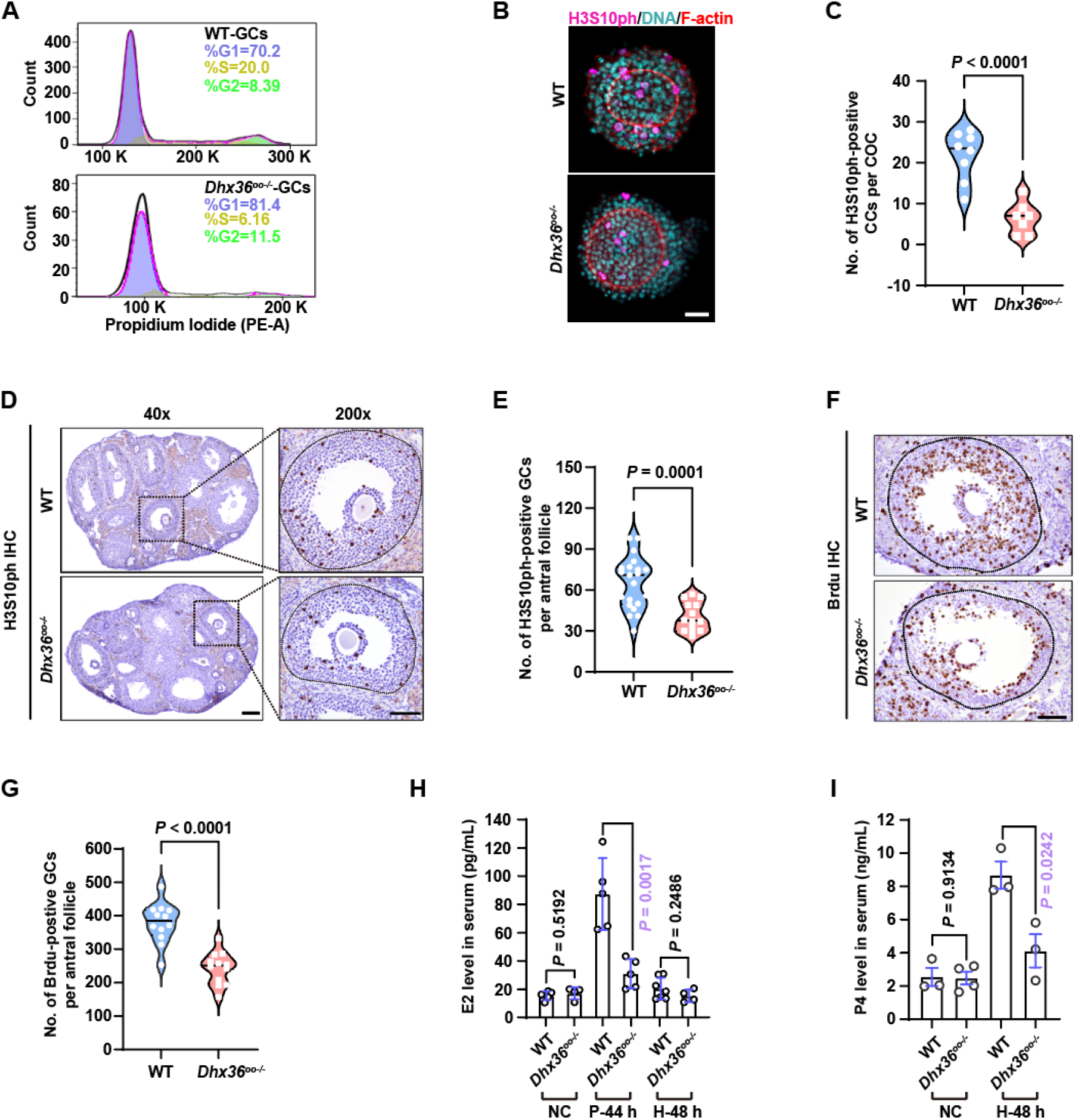
Oocyte-specific deletion of *Dhx36* suppresses cumulus and granulosa cell proliferation in response to hormonal stimulation. A: Flow cytometry histograms showing cell cycle distribution of granulosa cells isolated from antral follicles of WT and *Dhx36^oo-/-^*mice after 44 h of PMSG treatment. Cells were stained with propidium iodide. The percentages of cells in G0/G1, S, and G2/M phases are indicated. **B:** IF staining of phosphorylated histone H3 at serine 10 (H3S10ph) and F-actin in COCs isolated from 4-week-old WT and *Dhx36^oo-/-^* mice after treatment with PMSG for 44 h. (Scale bar: 20 μm). **C:** Quantification of H3S10ph-positive cumulus cells per COC. **D:** IHC staining of H3S10ph in ovarian sections from 4-week-old WT and *Dhx36^oo-/-^* mice after PMSG treatment for 44 h. (Scale bar: 100 μm and 50 μm, left to right). **E:** Quantification of H3S10ph-positive granulosa cells per antral follicle section. **F:** IHC staining of BrdU in ovarian sections from 4-week-old WT and *Dhx36^oo-/-^* mice after PMSG treatment for 44 h. (Scale bar: 50 μm). **G:** Quantification of BrdU-positive granulosa cells per antral follicle section. **H:** Serum estradiol (E2) level in WT and *Dhx36^oo-/-^* mice under conditions including untreated control (NC), 44 h after PMSG treatment (P-44 h), 48 h after hCG treatment following PMSG priming (H-48 h:). **I:** Serum progesterone (P4) level in WT and *Dhx36^oo-/-^* mice with or without hCG treatment. All quantitative data are presented as mean ± SEM. *P* values were calculated using unpaired *t*-tests.

### SMAD signaling is disrupted in cumulus cells following oocyte deprivation of *Dhx36*

To characterize the ovulatory defects, we next examined COC morphology at 2, 4, 8 and 16 h post-hCG injection following PMSG priming. Within WT antral follicles, cumulus cells began transitioning from a cuboidal to a polarized morphology by 4 h post-hCG, underwent robust expansion by hCG-8 h, and were accompanied by CL formation by 16 h (**Fig. 3A**). In contrast, cumulus cells within *Dhx36^oo-/-^* antral follicles exhibited insufficient polarization at 8 h post-hCG and became detached from oocytes by 16 h (**Fig. 3A**). Consistently, fewer COCs were observed from the oviductal ampulla of *Dhx36^oo-/-^* mice at 16 h post-hCG (**Fig. S2A**), and these COCs exhibited impaired expansion, characterized by reduced diameter and loss of the polarized corona radiata (**Fig. 3B**). *In vitro* epidermal growth factor (EGF) stimulation, which induces robust COC expansion through activation of the epidermal growth factor receptor (EGFR)–extracellular cellular signal-regulated kinase 1 and 2 (ERK1/2, also known as MAPK3/1) signaling pathway in cumulus cells^19^, failed to induce cumulus expansion in *Dhx36^oo-/-^* COCs, with disaggregated and detached cumulus cells observed after 16 h of EGF treatment (**Fig. 3C, D**). In addition, the expression of cyclooxygenase 2 (COX-2, also known as PTGS2), a key enzyme required for cumulus expansion^20,21^, was markedly reduced in *Dhx36^oo-/-^* ovaries at 8 h post-hCG (**Fig. 3E, F; Fig. S2B**).

**Figure 3.**
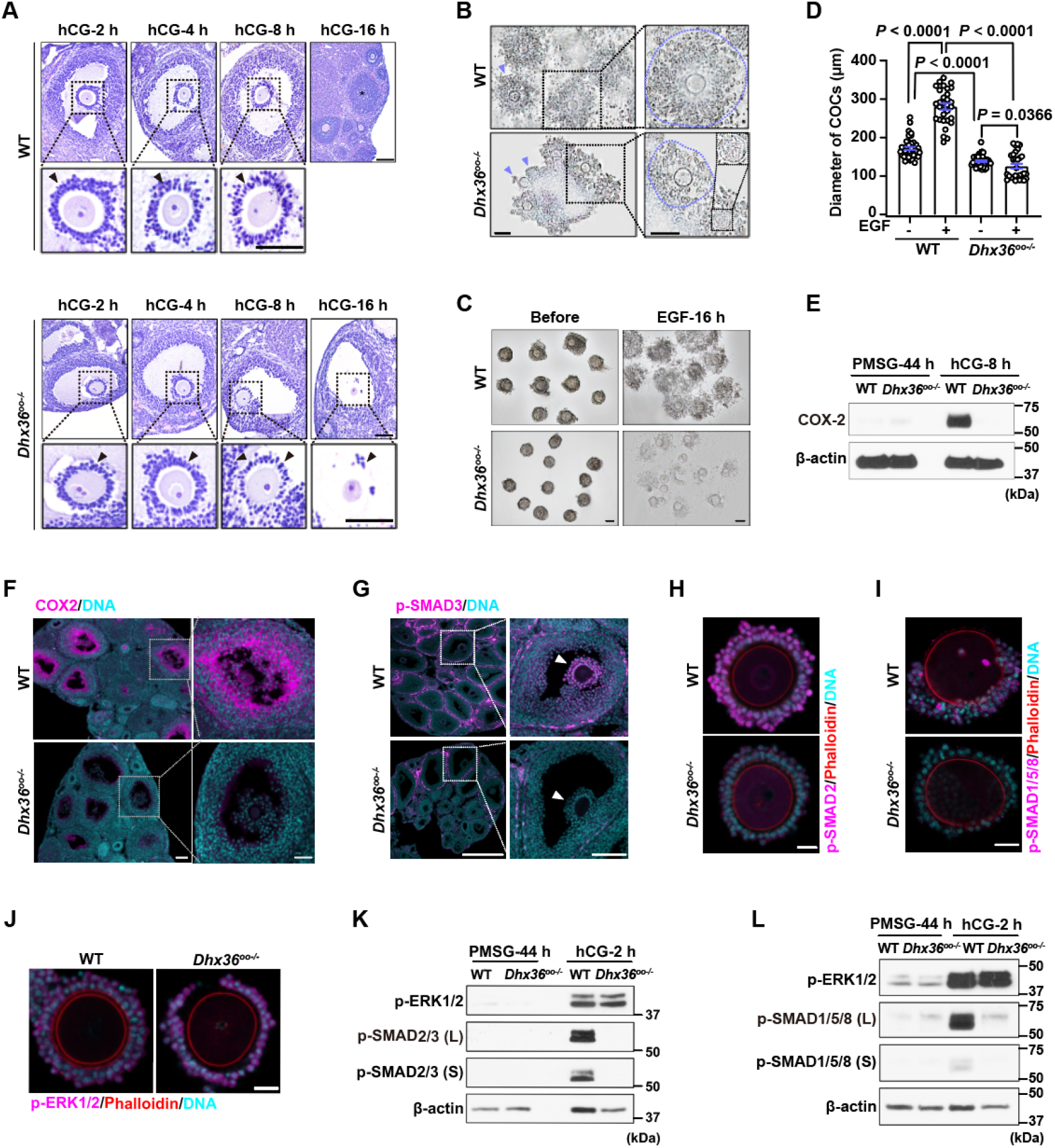
Oocyte-specific loss of *Dhx36* attenuates SMAD signaling pathway and cumulus expansion. A: H&E staining of ovarian sections and COCs from WT and *Dhx36^oo-/-^* mice at 2 h, 4 h, 8 h, and 16 h post-hCG injection following PMSG priming for 44 h. Black triangles indicate cumulus cells; asterisk indicates CL. (Scale bar: 100 μm). **B:** COCs collected from the oviductal ampulla of WT and *Dhx36^oo-/-^* mice treated with hCG for 16 h following PMSG priming for 44 h. Purple triangles indicate cumulus cells, and purple dashed circles denote individual COCs. (Scale bar: 100 μm). **C:** Bright-field images of COCs isolated from WT and *Dhx36^oo-/-^* mice before and after in vitro expansion induced by EGF (100 ng/mL) for 16 h. **D:** Quantification of COC diameter before and after EGF treatment. **E:** WB analysis of COX-2 protein level in ovarian lysates from WT and *Dhx36^oo-/-^*mice at 44 h post-PMSG (PMSG-44 h) and 8 h post-hCG following PMSG priming (hCG-8 h). **F:** IF staining of COX-2 in ovarian sections from WT and *Dhx36^oo-/-^*mice at 8 h post-hCG injection. (Scale bar: 100 μm). **G:** IF staining of p-SMAD3 in ovarian sections from WT and *Dhx36^oo-/-^* mice at 2 h post-hCG injection following PMSG priming. White triangles indicate cumulus cells. (Scale bar: 100 μm). **H–J:** IF images showing p-SMAD2 (H), p-SMAD1/5/8 (I), and p-ERK1/2 (J) signal in COCs collected from 4-week-old WT and *Dhx36^oo-/-^* mice at 2 h post-hCG injection after PMSG priming. (Scale bar: 20 μm). **K–L:** WB analysis of p-ERK1/2, p-SMAD1/5/8 and p-SMAD2/3 protein level in ovarian lysates from 4-week-old WT and *Dhx36^oo-/-^* mice at 44 h post-PMSG (PMSG-44 h) and 2 h post-hCG injection following PMSG priming (hCG-8 h). All quantitative data are presented as mean ± SEM. *P* values were calculated by unpaired *t*-tests.

Furthermore, the phosphorylation level of SMAD2/3 and SMAD1/5/8 were significantly reduced in cumulus cells from antral follicles of *Dhx36^oo-/-^* mice at 2 h post-hCG (**Fig. 3G–I; Fig. S2C–G, Fig. 3K, L**). In contrast, ERK1/2 phosphorylation was normally induced in cumulus cells from antral follicles of *Dhx36^oo-/-^* mice (**Fig. 3J–L; Fig. S2H**). Collectively, these results indicate that oocyte-specific loss of *Dhx36* impairs cumulus expansion and selectively disrupts TGF-β superfamily-dependent SMAD signaling in cumulus cells, while leaving ERK1/2 activation largely intact.

### Oocyte-specific *Dhx36* loss impairs cumulus cell function and adhesion

To assess the functional properties of cumulus cells, we performed COC reconstitution assays by exchanging cumulus cells between WT and *Dhx36^oo-/-^* mice. After excluding dead oocytes, reconstitution efficiency, defined as the proportion of successfully reconstituted COCs among all viable oocytes, including denuded oocytes, was reduced by approximately 50% in the *Dhx36^oo-/-^*-derived cumulus cells group compared with the WT-derived group, which showed an efficiency of over 80% (**Fig. 4A; Fig. S3A, B**). Furthermore, cumulus cell detachment was frequently observed during *in vitro* EGF-induced cumulus expansion in reconstituted COCs containing *Dhx36^oo-/-^*-derived cumulus cells (**Fig. 4B**), indicating impaired adhesive and expansion capabilities of *Dhx36^oo-/-^*-derived cumulus cells.

**Figure 4.**
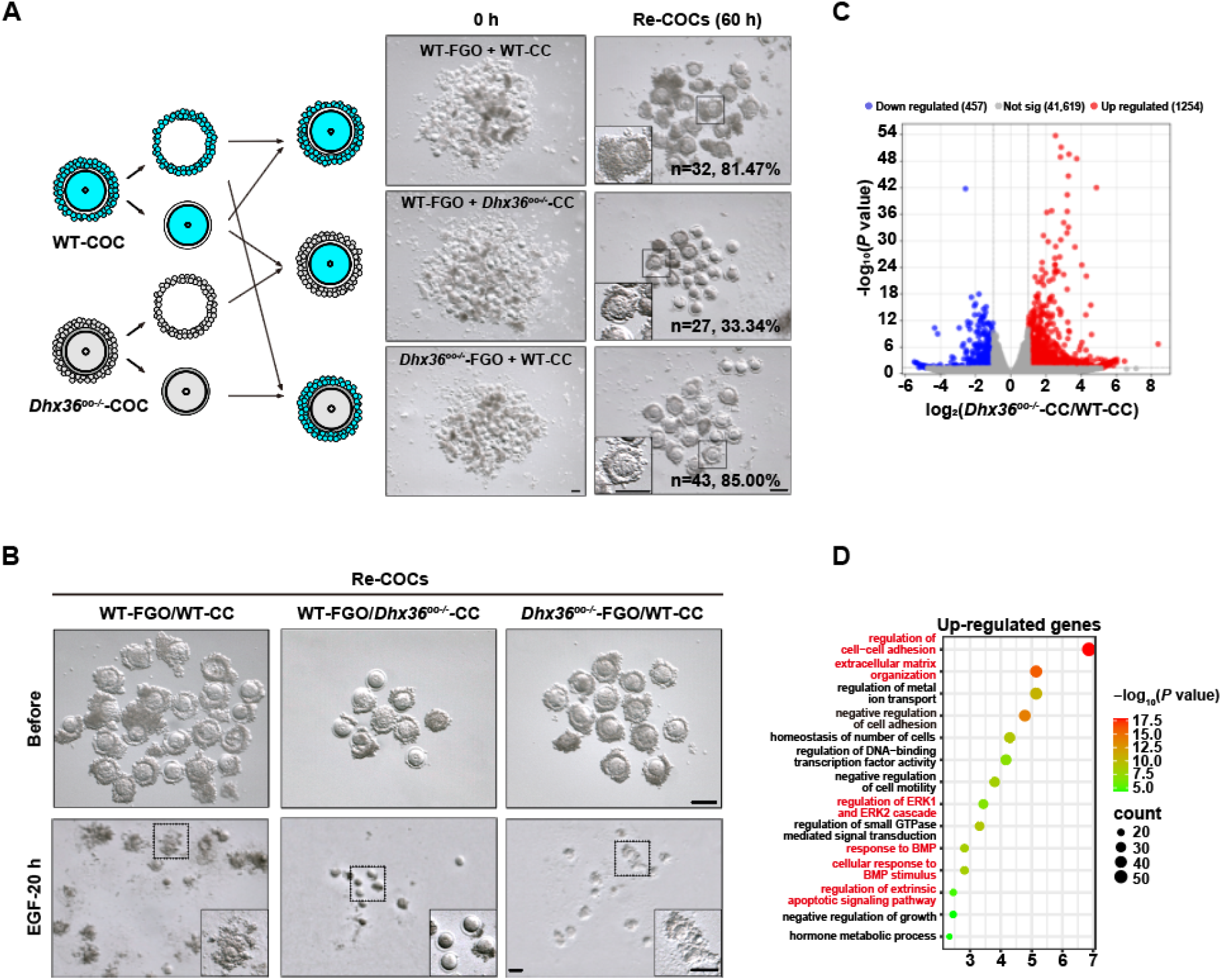
Disrupted gene expression and defective adhesion in cumulus cells surrounding *Dhx36*-deficient oocytes. A: Schematic diagram of COC reconstitution strategy and bright-field images of reconstituted COCs. (Scale bar: 100 μm). The total number of used oocytes (n) and reconstitution success rates are indicated at the bottom right of the images. COCs from WT and *Dhx36^oo-/-^*mice were dissociated into FGO (designated WT-FGO and *Dhx36^oo-/-^*-FGO) and cumulus cells (designated WT-CC and *Dhx36^oo-/-^*-CC), which were then were reconstituted. Cumulus cells from WT mice (WT-CC) were reconstituted with WT FGO (WT-FGO) as a control. **B:** Bright-field images showing cumulus expansion of reconstituted COCs in all groups before and 20 h after EGF induction. (Scale bar: 100 μm). **C:** Volcano plot showing differentially expressed genes (DEGs) (|log2FC| ≥ 1, *P* value < 0.05) in cumulus cells from *Dhx36^oo-/-^* (*Dhx36^oo-/-^*-CCs) versus WT (WT-CCs) mice. **D:** Gene ontology enrichment analysis of biological processes among upregulated genes in *Dhx36^oo-/-^* cumulus cells.

To further define the molecular consequences of impaired cumulus cell function, we performed RNA-seq on cumulus cells isolated from WT and *Dhx36^oo–/–^* mice. Principal component analysis (PCA) revealed clear separation between WT and *Dhx36^oo–/–^* cumulus cells (**Fig S3C**). Differential expression analysis identified 1254 upregulated and 457 downregulated genes in *Dhx36^oo-/-^*-derived cumulus cells compared to WT-derived cumulus cells (**Table S4; Fig. 4C**). Interestingly, gene ontology analysis revealed that upregulated genes in *Dhx36^oo-/-^*-derived cumulus cells were significantly enriched in cell–cell adhesion and extracellular matrix organization, suggesting a potential compensatory response to impaired cumulus cell adhesion (**Fig. 4D**). Downregulated genes were enriched in biological processes related to cellular cation homeostasis and regulation of transmembrane transporter activity (**Fig. S3D**). Together, these findings demonstrate that oocyte-specific loss of *Dhx36* induces persistent alterations in gene expression in cumulus cells and is associated with impaired cumulus cell adhesion and expansion.

### Defective BMPs delivery caused by aberrant membrane architecture in *Dhx36*-deficient oocytes

Since the defects in cumulus cells potentially result from impaired OSF delivery and oocyte-cumulus intracellular communication, we next examined OSF expression and the relevant structural features in both oocytes and cumulus cells. Notably, the expression level of GDF-9 and BMP-15 were comparable between FGO form WT and *Dhx36^oo-/-^* mice, whereas phosphorylation of ERM (ezrin/radixin/moesin), which is associated with microvilli organization, was significantly reduced in *Dhx36^oo-/-^* oocytes (**Fig. 5A**). Consistently, IF and transmission electron microscopy (TEM) revealed markedly reduced microvilli density in *Dhx36^oo-/-^* oocytes (**Fig. 5B–E**). Phalloidin staining of F-actin and TEM also demonstrated reduced TZP density in *Dhx36^oo-/-^* oocytes (**Fig. 5F–I**). Accordingly, GDF-9 enrichment within the TZP region was significantly reduced in *Dhx36^oo-/-^* oocytes, accompanied by a more diffuse and disorganized distribution throughout the oocyte cytoplasm (**Fig. 5J**), suggesting impaired trafficking of GDF-9 toward cumulus cells. Similarly, BMP-15 signal intensity within the TZP region was reduced in *Dhx36^oo-/-^* oocytes (**Fig. 5K**). Consistent with impaired intercellular communication, Lucifer Yellow microinjection and diffusion assays further revealed reduced transfer from oocytes to cumulus cells (**Fig. 5L–N**).

**Figure 5.**
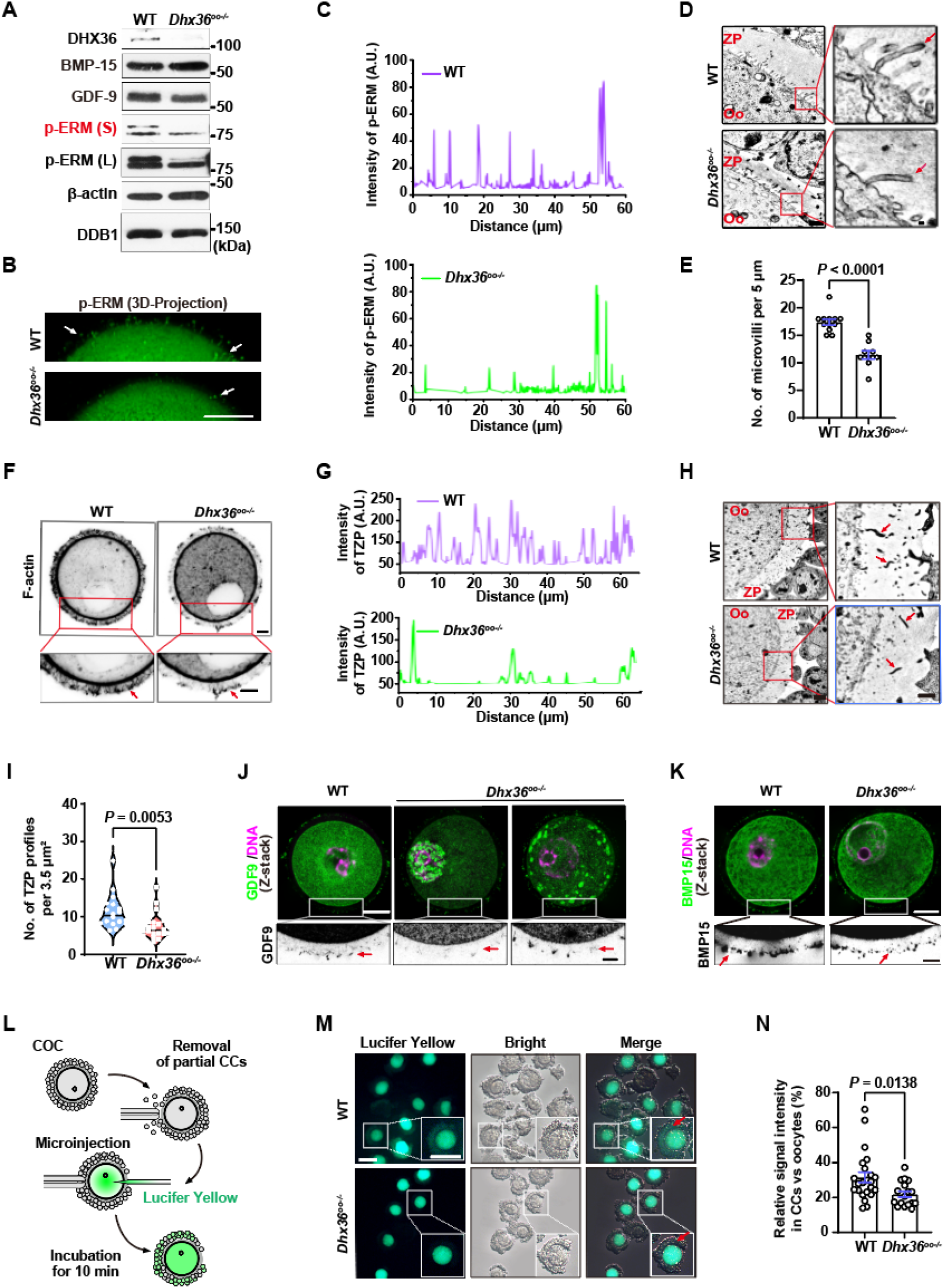
Impaired microvilli compromise the delivery of OSFs from *Dhx36-*null oocytes to cumulus cells. A: Western blot showing protein level of DHX36, BMP-15, GDF-9, and p-ERM in FGO from FGO from WT and *Dhx36^oo-/-^* mice. β-actin and DDB1 were used as internal controls. p-ERM expression was detected under short exposure (S) and long exposure (L). **B:** IF staining of p-ERM in WT and *Dhx36^oo-/-^* FGO. (Scale bar: 20 μm). Z-stack images were merged using the Auto-Blend Layers function in Photoshop. **C:** Distribution of p-ERM fluorescence intensity along the microvilli-enriched region of the oocyte cortex shown in (B). **D:** Transmission electron microscopy (TEM) images showing microvilli structure of oocytes within secondary follicles from WT and *Dhx36^oo-/-^* mice. ZP, zonal pellucida; Oo, oocyte. Red arrows indicate individual microvilli protrusion. (Scale bars: 1 μm, 200 nm, 100 nm, from left to right). **E:** Quantification of microvilli density by measuring number of microvilli protrusion per 5 μm based on images in (D). **F:** Inverted fluorescence images of F-actin labeled with phalloidin in WT and *Dhx36^oo-/-^* FGO. Red arrows indicate transzonal projections (TZPs). (Scale bar: 10 μm). **G:** Quantification of TZP fluorescence intensity along straightened images of the zona pellucida in (F). **H:** TEM images of TZPs within zona pellucida in COCs from WT and *Dhx36^oo-/-^* mice. Red arrows indicate truncated TZPs. (Scale bars: 2 μm, 1 μm, from left to right). **I:** Quantification of TZP number of TZP per 3.5 μm² based on TEM images in (H). **J, K:** Immunofluorescence staining of BMP-15 (J) and GDF-9 (K) in WT and *Dhx36^oo-/-^* FGO stripped from COCs. Red arrows denote BMP-15 and GDF-9 signal within TZPs. (Scale bars: 20 μm, 5 μm, from up to down). **L:** Schematic diagram of lucifer yellow microinjection assay. Cumulus cells were partially removed to enable subsequent microinjection lucifer yellow into oocytes. **M:** Fluorescence images of COCs incubated for 10 min following lucifer yellow microinjection. (Scale bar: 100 μm). **N:** Quantification of lucifer yellow signal intensity in cumulus cells relative to oocytes. Data in histogram and violin plot are presented as mean ± SEM. *P* values were calculated by unpaired *t*-tests.

### DHX36 safeguards autophagy and organelle homeostasis in oocytes

To investigate the molecular basis of the defects in *Dhx36^oo-/-^* oocytes, we performed low-input proteomics on FGO from WT and *Dhx36^oo-/-^* mice. More than 5,000 proteins were identified in each group (**Table S5**), with 614 upregulated and 628 downregulated proteins in *Dhx36^oo-/-^* FGO (**Fig. 6A; Fig. S4A–C**). Downregulated proteins were enriched in autophagy, phospholipid metabolism, organelle fusion, and mitophagy, whereas upregulated proteins were mainly associated with RNA splicing and ribonucleoprotein complex biogenesis (**Fig. 6B; Fig. S4D–E**). Among the downregulated autophagy-related proteins were key regulators of autophagosome biogenesis, vesicle fusion, and selective autophagy, including MAPK3/1 (ERK1/2)^22^, FOXO3^23–27^, UVRAG and ATG2A^28,29^, and multiple SNARE proteins (SNAP47, SNAP29, STX16, VAMP3, and VTI1B)^30–35^ (**Fig. 6C**).

**Figure 6.**
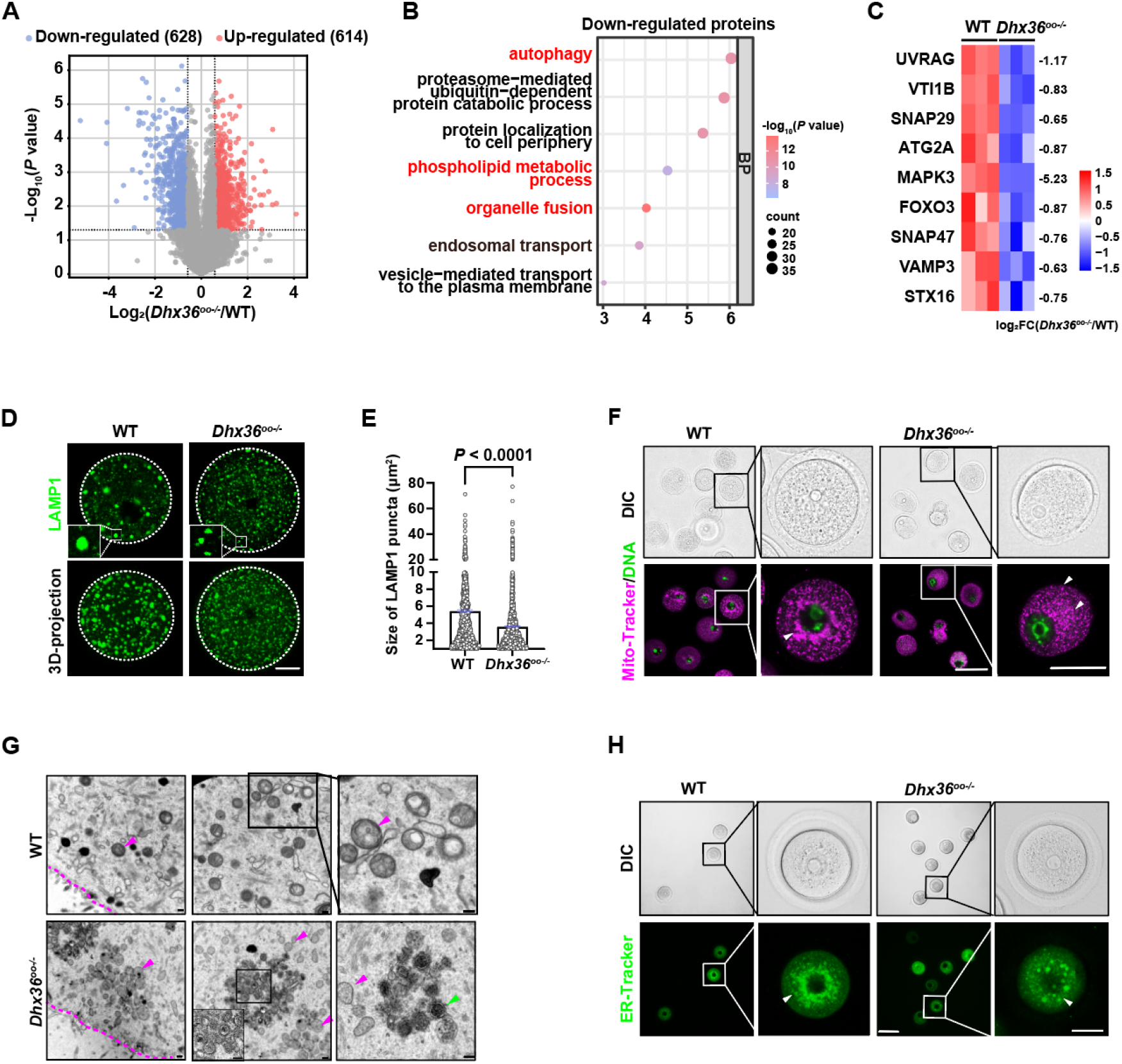
Autophagy defects accompanied by aberrant lysosome, mitochondria and endoplasmic reticulum morphology in *Dhx36*-deficient oocytes. A: Volcano plot of differentially expressed proteins in FGO from WT and *Dhx36^oo-/-^* mice. The cutoff criteria are |log₂FC| ≥ 0.585 (equivalent to ≥ 1.5-fold change) and *P* value < 0.05). **B:** Gene ontology enrichment analysis of downregulated proteins in *Dhx36^oo-/-^* FGO. **C:** Heatmap showing the expression level of key autophagy-related proteins in WT and *Dhx36^oo-/-^* FGO. **D:** Immunofluorescence staining of LAMP1 in WT and *Dhx36^oo-/-^*FGO. (Scale bar: 20 μm). **E:** Quantification of LAMP1-positive puncta size in WT and *Dhx36^oo-/-^* FGO. Data is presented as mean ± SEM. *P* value was calculated using unpaired *t*-test. **F:** Live-cell imaging of mitochondria labeled with Mito-Tracker Red in WT and *Dhx36^oo-/-^*FGO. White triangles indicate mitochondrial aggregates. (Scale bar: 100 μm). **G:** TEM images showing mitochondrial morphology in oocytes from WT and *Dhx36^oo-/-^* mice. (Scale bar: 200 nm). Pink dashed lines outline the oocyte boundaries; pink triangles indicate mitochondria; green triangles indicate aberrantly accumulated mitochondria that may represent intermediate structures undergoing autophagic degradation in *Dhx36^oo-/-^* oocytes. In *Dhx36^oo-/-^*oocytes, aberrant mitochondria (100–450 nm in diameter) exhibited a shrunken morphology and disrupted cristae and were predominantly localized at the oocyte periphery, whereas WT oocytes contained oval, double-membraned mitochondria (300–500 nm in diameter). **H:** Live-cell imaging of the ER labeled with ER-Tracker Green in WT and *Dhx36^oo-/-^*FGO. (Scale bars: 100 μm, 20 μm, from left to right).

Consistent with these proteomic changes, *Dhx36^oo-/-^*oocytes exhibited reduced LAMP1-positive puncta (**Fig. 6D–E**). Mitochondria lost their normal perinuclear clustering and displayed abnormal morphology and disrupted cristae (**Fig. 6F–G**), while the ER became fragmented and disorganized compared with its organized perinuclear distribution in WT oocytes (**Fig. 6H**). Together, these findings indicate that DHX36 deficiency disrupts lysosomal, mitochondrial, and ER homeostasis, consistent with impaired autophagy and organelle turnover in *Dhx36^oo-/-^* oocytes.

### DHX36 sustains phospholipid metabolism and homeostasis in oocytes

Proteomic analysis revealed broad downregulation of proteins involved in lipid metabolism, including PCYT1B, which catalyzes the rate-limiting step of phosphatidylcholine synthesis^36^; GPAT2, involved in glycerolipid synthesis^37^; LPIN1 and LPIN2, regulators of triacylglycerol (TAG) synthesis and phospholipid homeostasis^38,39^; and the lipid catabolic enzymes LPL and ABHD6^40,41^ (**Fig. S5A**). Consistent with these changes, *Dhx36^oo-/-^* FGO exhibited smaller and more dispersed Nile Red-positive lipid droplets compared with the consolidated droplets in WT oocytes (**Fig. 7A**). Low-input LC-MS profiling further revealed altered lipid homeostasis (**Table S6**). Neutral lipid profiles showed limited separation between genotypes, with comparable diacylglycerol (DAG) levels but significantly increased TAG in *Dhx36^oo-/-^* oocytes (**Fig. S5B; Fig. 7B**). In contrast, phospholipid profiles clearly separated WT and *Dhx36^oo-/-^*FGO (**Fig. 7C**), with significantly reduced lysophosphatidylcholine (LPC) and lysophosphatidylethanolamine (LPE) level (**Fig. 7D**). Among 87 detected phospholipids, 21 were downregulated and 3 were upregulated in *Dhx36^oo-/-^*FGO, with most differential species belonging to the phosphatidylcholine (PC), phosphatidylethanolamine (PE), and LPE classes (**Fig. 7E–G; Fig. S5C**). Together, these findings demonstrate disrupted neutral lipid and phospholipid homeostasis in *Dhx36^oo-/-^* oocytes, consistent with the membrane and organelle abnormalities observed above.

**Figure 7.**
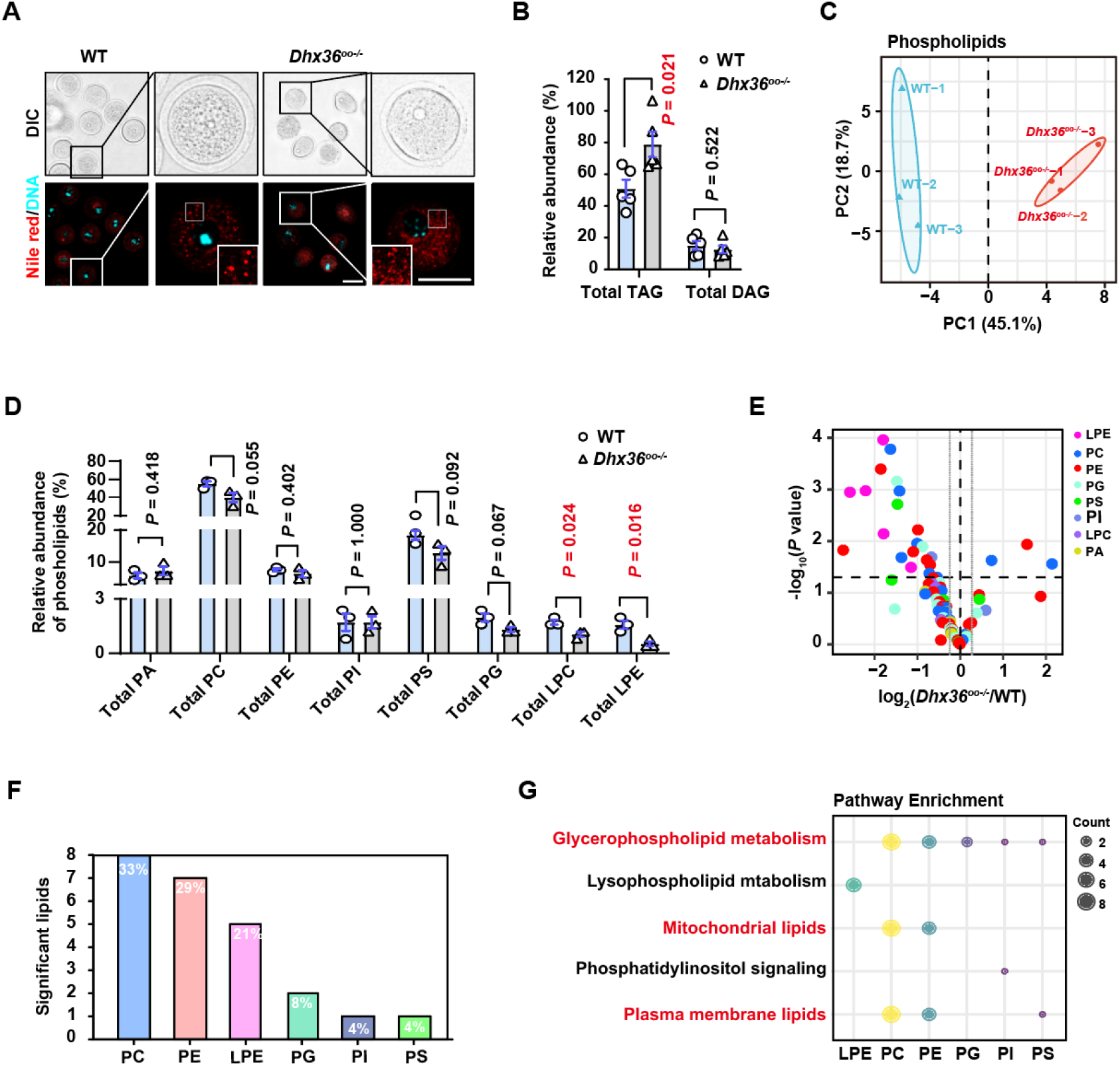
Dysregulation of lipid metabolism in *Dhx36^oo-/-^* oocytes. A: Live-cell imaging of neutral lipids stained with Nile Red in WT and *Dhx36^oo-/-^* FGO. (Scale bar: 100 μm). **B:** Abundance of total triacylglycerol (TAG) and diacylglycerol (DAG) in WT and *Dhx36^oo-/-^* FGO. **C:** PCA of phospholipid profiles in FGO from WT and *Dhx36^oo-/-^*mice. **D:** Abundance of different phospholipid classes in WT and *Dhx36^oo-/-^*FGO, including phosphatidic acid (PA), phosphatidylcholine (PC), phosphatidylethanolamine (PE), phosphatidylinositol (PI), phosphatidylserine (PS), phosphatidylglycerol (PG), lysophosphatidylcholine (LPC), and lysophosphatidylethanolamine (LPE). **E:** Volcano plot showing differentially abundant phospholipids in *Dhx36^oo-/-^* versus WT FGO. Vertical grey lines indicate a fold change threshold of ≥ 1.2 (|log₂(fold change)| ≥ 0.263); the horizontal black dashed line indicates *P* < 0.05 (-log₁₀(*P*) ≥ 1.30). **F:** Proportion of differentially abundant phospholipid classes. **G:** Bubble plot showing pathways enriched in differentially abundant phospholipids. Quantitative data are presented as mean ± SEM. *P* values were calculated using unpaired *t*-tests.

### Downregulated genes in *Dhx36*-null oocytes are enriched with G-quadruplex motifs within their promoter regions

Given the role of DHX36 in chromatin architecture and transcription regulation^42^, we hypothesized that the defects in membrane assembly, autophagy, organelle homeostasis, and lipid metabolism arise from transcriptional dysregulation in *Dhx36^oo-/-^*oocytes. Consistently, mRNA levels and protein abundance showed a strong positive correlation (**Fig. 8A**), and 232 genes were concordantly downregulated at both levels using a 1.5-fold change threshold (|log₂FC| ≥ 0.585) (**Fig. 8B**). These genes included regulators of organelle function, lipid metabolism, membrane assembly, and autophagy (**Fig. 8C**), and exhibited reduced chromatin accessibility at transcription start sites^42^ (**Fig. 8D; Fig. S6A–C**).

**Figure 8.**
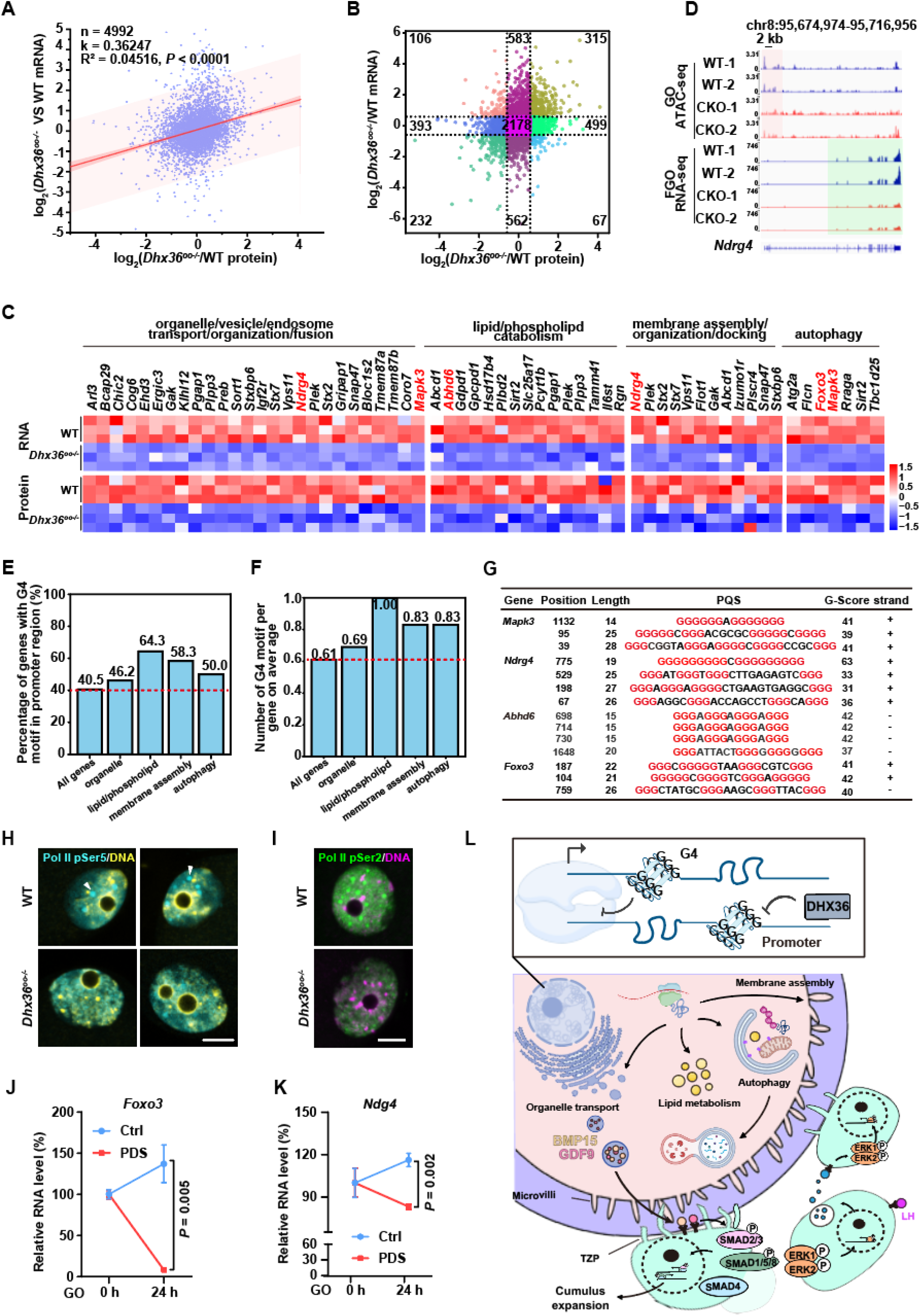
DHX36 safeguards transcription of genes involved in key pathways. A: Scatter plot illustrating the correlation between mRNA and protein expression level in *Dhx36^oo-/-^* versus WT FGO. Red line represents the linear fit. **B:** Classification of genes with differentially expressed RNA and protein level, divided into nine regions based on scatter plot. Lines indicate a 1.5-fold change threshold (|log₂FC| ≥ 0.585). **C:** Heatmap showing concordant downregulation of both RNA and protein down-regulated expression of keys genes involved in gene ontology pathways, including organelle/vesicle transport or fusion, lipid/phospholipid metabolism, membrane assembly, and autophagy in *Dhx36^oo-/-^*oocytes compared to WT controls. Three key genes, *Ndrg4, Mapk3* and *Foxo3* are highlighted in red. **D:** Representative Integrative Genomics Viewer (IGV) tracks showing ATAC-seq and RNA-seq data for *Ndrg4* in growing oocytes (GO) and FGO from WT and *Dhx36^oo-/-^*mice. **E:** Histogram showing the percentage of gene containing putative G-quadruplex (PQS) motifs within their promoter regions for the gene sets from the enriched pathways shown in (C). The set of all genes identified in both RNA-seq and proteomic data was used as a control. **F:** Histogram showing the average number of PQS motifs per gene within the promoter region for the gene sets shown in (C). **G:** Position, length, strand and G-score of PQS motifs identified in the promoter regions of *Mapk3*, *Ndrg4*, and *Foxo3*. The position indicates the starting nucleotide number of each G4 motif within the 2 kb promoter sequence. The strand indicates the DNA strand as defined by UCSC Genome Browser. **H:** Immunofluorescence staining of RNA polymerase II Ser5 phosphorylation (Pol II pSer5) in growing oocytes from WT and *Dhx36^oo-/-^*mice. White triangles indicate the Pol II pSer5 foci in oocyte nucleus. (Scale bar: 20 μm). **I:** Immunofluorescence staining of RNA polymerase II Ser2 phosphorylation (Pol II pSer2) in growing oocytes from WT and *Dhx36^oo-/-^* mice. (Scale bar: 20 μm). **J, K:** RNA level of *Foxo3* (J) and *Ndrg4* (K) in growing oocytes before and after PDS treatment for 24 h, as measured by RT-qPCR. Data are presented as mean ± SEM. *P* values were calculated by unpaired *t*-tests. **L:** Schematic diagram illustrating the influence and potential mechanism of DHX36 on biological processes in oocytes and its subsequent effects on signaling pathways and cumulus expansion in surrounding cumulus cells. Luteinizing hormone, LH.

Because G4s are enriched in open chromatin^43^ and promoter regions, with approximately 40% of genes containing at least one promoter G4 motif^44^, we examined promoter G4 motifs using QGRS Mapper^45^. Pathway genes showed a higher proportion of strict G4 motifs (46.2–64.3%) than all genes (40.5%), as well as more G4 motifs per gene (0.69–1.0 vs. 0.61) (**Table S3; Fig. 8E–F**). Several key downregulated genes, including *Ndrg4, Mapk3, Abhd6,* and *Foxo3*, contained high-scoring promoter G4 motifs (**Fig. 8C, G**), with three identical G4 motifs identified in *Abhd6*. Synthetic oligonucleotides corresponding to these motifs showed strong G4-forming propensity in vitro using the CYTO probe (**Fig. S6D–I**), supporting their potential to form G4 structures.

Consistent with impaired transcription, RNA Polymerase II Ser5 and Ser2 phosphorylation, markers of transcription initiation and elongation, respectively, were dispersed or markedly reduced in *Dhx36^oo-/-^* growing oocytes (**Fig. 8H, I; Fig. S6J, K**), resembling the effect of the G4 stabilizer pyridostatin (PDS) (**Fig. S6L–O**). PDS treatment reduced *Foxo3* and *Ndrg4* mRNA levels compared with control oocytes, whereas *Mapk3* was unexpectedly increased (**Fig. 8J, K; Fig. S6P**), indicating that G4 stabilization can exert gene-specific effects and does not fully phenocopy DHX36 depletion.

Together, these findings link DHX36 deficiency to reduced expression of genes involved in membrane assembly, organelle homeostasis, lipid metabolism, and autophagy, whose promoters are enriched in G4 motifs. We propose that unresolved promoter G4s may contribute to impaired RNA Polymerase II activity and transcription of these genes, thereby disrupting oocyte structural homeostasis and OSF delivery and ultimately contributing to defective ovulation (**Fig. 8L**).

## Discussion

This study identifies the G4 helicase DHX36 as an oocyte-intrinsic factor required for ovulation and female fertility by maintaining oocyte structural homeostasis and supporting OSF delivery to surrounding cumulus cells. The severe cumulus expansion defect in *Dhx36^oo-/-^* mice, together with impaired SMAD2/3 and SMAD1/5/8 activation despite normal ERK1/2 activation, indicates that DHX36 deficiency primarily disrupts the oocyte-derived OSF–SMAD signaling axis rather than the LH/EGF–ERK pathway^46–48^ (**Fig. 8L**). Because GDF-9 and BMP-15 are major oocyte-derived signals required for SMAD activation in cumulus cells, these findings support defective OSF delivery as a major cause of impaired cumulus expansion. Moreover, the failure of *Dhx36^oo-/-^*cumulus cells to efficiently reconstitute COCs with WT oocytes suggests that prolonged impairment of oocyte-derived signaling during follicle growth may cause persistent defects in cumulus cell–oocyte communication, potentially involving defective TZP development and maintenance. Interestingly, the upregulation of adhesion-and BMP/TGF-β-related genes despite impaired adhesion and signaling may represent a compensatory response to chronic insufficiency of oocyte-derived signals.

Our findings further suggest that impaired OSF delivery is associated with broad disruption of oocyte membrane and organelle homeostasis. DHX36 deficiency affects microvilli/TZP-associated structures, phospholipid metabolism, vesicle and lysosome homeostasis, and autophagy, which may collectively compromise the membrane remodeling and vesicular processes required for efficient OSF transport. Phospholipids are fundamental components of cellular membranes and are essential for membrane organization, vesicle trafficking, and organelle function^49^. These defects may therefore reinforce one another, impairing membrane integrity and organelle turnover and ultimately disrupting the delivery of oocyte-derived signals to cumulus cells. Thus, DHX36 appears to support oocyte–cumulus communication not through a single structural component, but by maintaining the broader membrane and organelle homeostasis of the oocyte.

At the molecular level, integrated transcriptomic and proteomic analyses revealed coordinated downregulation of genes involved in membrane assembly, phospholipid metabolism, organelle homeostasis, and autophagy. Notably, these affected genes exhibited a higher prevalence of potential promoter G4-forming sequences than the genome-wide background, suggesting that they may represent a class of genes sensitive to DHX36-dependent G4 regulation. Although direct binding of DHX36 to these promoter G4 structures remains to be established, the observed G4 enrichment, together with the effects of G4 stabilization on the expression of selected genes and RNA polymerase II activity, supports a potential role for promoter G4 dynamics in the transcriptional regulation of these pathways. These findings suggest a model in which DHX36-dependent G4 regulation contributes to the expression of genes required for membrane and organelle homeostasis, thereby linking oocyte-intrinsic molecular regulation to OSF delivery, oocyte–cumulus communication, and ovulation (**Fig. 8L**).

## Data availability

The RNA-seq data generated in this study have been deposited in the Gene Expression Omnibus (GEO) database under accession number GSE333282. The mass spectrometry proteomics data have been deposited to the ProteomeXchange Consortium via the PRIDE partner repository with the dataset identifier PXD078772.

## Author Contributions

H.Y.F. conceived and supervised the study. Y.X.J., G.W.B., Y.L.C., K.Z., F.Y.S., B.Y.G., H.T.D., Y.Z.S., H.Y.S., S.Y.L., and Y.J.L. performed the experiments and conducted the methodology. Y.X.J. wrote the manuscript. Q.M.S., X.C., H.W., and C.Y. provided critical resources. H.Y.F. acquired the funding.

## Acknowledgements

We thank Wei-Na Shang and Zi-Yi Kang at the core facilities of the Life Sciences Institute Zhejiang University for support with TEM and confocal imaging. This work was supported by the National Key Research and Development Program of China (2024YFA1803000), the National Natural Science Foundation of China (U25A20658).

## Materials and Methods

### Animal Experimentation

*Dhx36^fl/fl^* female mice were crossed with *Dhx36^fl/fl^; Zp3-Cre* males to generate *Dhx36^fl/fl^; Zp3-Cre* female mice (*Dhx36^oo-/-^*), with Cre-negative littermate *Dhx36fl/fl* females used as WT controls. All animal procedures were approved by the Institutional Animal Care and Research Committee of Zhejiang University (Protocol No. ZJU20210309) and conducted in accordance with institutional guidelines and the ARRIVE guidelines.

### Oocyte collection

Growing oocytes were collected from 2-week-old female mice. FGO were isolated from 4–6-week-old mice at 44–48 h after PMSG injection. All oocytes were collected and cultured in M2 medium supplemented with 5 μM milrinone to maintain meiotic arrest.

### COC collection and cumulus expansion assay

Female mice (4 weeks old) were euthanized at 44–48 h after PMSG injection. COCs were collected from antral follicles by puncturing with a needle. The COCs were then cultured for 16–20 h in Dulbecco’s Modified Eagle Medium (DMEM, Gibco, D9800-13) supplemented with 20 mM HEPES, 0.3% BSA, 1% fetal bovine serum (FBS), 1% penicillin-streptomycin (P/S), and 100 ng/mL EGF. After EGF treatment for 16 h, COCs were imaged using a microscope.

### Hematoxylin and eosin (H&E) staining and ovarian follicle counting

Ovaries were collected, rinsed with PBS, fixed overnight in 10% formaldehyde at 4°C, and processed for paraffin embedding. Serial sections (6–7 μm) were prepared and subjected to H&E staining following standard procedures. Ovarian follicles were classified and counted based on standard morphological criteria: primordial follicles contained an oocyte surrounded by a single layer of flattened granulosa cells; primary follicles contained a single layer of cuboidal granulosa cells; secondary follicles had multiple granulosa cell layers without an antrum; and antral follicles contained multiple granulosa cell layers with one or more antral spaces. Only oocytes with a clearly visible nucleus were counted to avoid duplicate counting. For granulosa cell quantification, antral follicle images were analyzed using ImageJ. Nuclear regions were segmented by thresholding and the watershed algorithm was applied to separate adjacent nuclei. Granulosa cells within manually defined regions of interest were subsequently counted.

### Follicle number counting

Follicles in H&E-stained ovarian sections were classified: primary (oocyte with one layer of cuboidal granulosa cells), early secondary (2–3 granulosa cell layers), late secondary (>3 layers, no antrum), antral (visible antrum), and late antral (distinct cumulus and mural granulosa cell layers). Only follicles with an oocyte exhibiting a clearly visible nucleus were counted.

### Low-input proteomics

Low-input proteomics was performed as previously described53 with modifications. FGO from WT and *Dhx36^oo-/-^* mice were treated with acidic Tyrode’s solution to remove the zona pellucida. After washing with PBS, 20 zona-free oocytes per group were collected in protein LoBind tubes and stored at −80°C. Samples were lysed in 1% sodium deoxycholate in 20 mM Tris-HCl (pH 8.5), heated at 100°C for 10 min, and sonicated. Proteins were digested with trypsin and LysC (1:100, w/w), and peptides were desalted using SDB-RPS StageTips, dried, and reconstituted in 0.1% formic acid. Peptides were analyzed by LC-MS/MS using a Vanquish Neo UHPLC system coupled to an Orbitrap Astral mass spectrometer. Data were acquired in data-independent acquisition (DIA) mode. Raw data were processed using Spectronaut v18 with the directDIA workflow against the reviewed mouse UniProt database. Proteins were identified and quantified with a 1% false discovery rate (FDR) at the PSM, peptide, and protein group levels.

### IHC and analysis

Paraffin-embedded ovarian sections were rehydrated and subjected to antigen retrieval in 10 mM sodium citrate buffer (pH 6.0) at 95°C for 15 min after quenching endogenous peroxidase with 3% H₂O₂. Sections were blocked with 5% goat serum and incubated overnight at 4°C with anti-H3ph10 antibody (CST, 9701S). Signals were detected using a biotin-conjugated secondary antibody, ABC complex, and DAB substrate (Vector kits PK-6100 and SK-4100), followed by hematoxylin counterstaining. H3ph10-positive granulosa cells were quantified using ImageJ after H-DAB color deconvolution and thresholding, with watershed and “Analyze Particles” applied to separate and count positive nuclei within defined regions of interest.

### BrdU staining and analysis

Female mice were intraperitoneally injected with Brdu (50 μg per gram of body weight) dissolved in PBS at 42–44 h after PMSG administration. 2 h after Brdu injection, ovarian tissues were collected and processed into formalin-fixed, paraffin-embedded sections. Sections were deparaffinized and rehydrated as described in the IHC procedure, rinsed with PBS, and then treated with 3% H₂O₂ for 10 min at 37 °C to block endogenous peroxidase activity. After washing with PBS, sections were subjected to DNA denaturation with 2 N HCl for 20 min at 37 °C, followed by incubation with 0.1% (w/v) trypsin in PBS for 30 min at 37 °C. After three PBS washes, sections were blocked with 5% goat serum for 1 h at room temperature and incubated with an anti-BrdU primary antibody (Sigma-Aldrich, B2531) at 4 °C overnight. Subsequent steps, including incubation with a biotinylated secondary antibody and DAB color development, were performed as described in the IHC section. The number of Brdu-positive granulosa cells was quantified using the same image analysis method outlined in the IHC analysis.

### Cell cytometry and cell cycle analysis of granulosa cells

For each group, three 3–4-week-old female mice were treated with PMSG and sacrificed 44 h later. Ovaries were collected, cleaned of surrounding tissues, and follicles were punctured with a 27-gauge needle in minimal PBS to release granulosa cells. Cell aggregates were dissociated with trypsin at 37°C for 30 min, and digestion was terminated with DMEM containing FBS. After removal of tissue debris, cells were collected by centrifugation, washed, and fixed in 70% ethanol under continuous vortexing for 30 min on ice, followed by storage at 4°C. For cell-cycle analysis, fixed cells were stained with propidium iodide (10 µg/mL) and RNase A (0.25 ng/mL) in PBS containing 1% FBS at 37°C for 30 min. Samples were filtered through a 70-µm cell strainer and analyzed using a CytoFLEX S flow cytometer. Single cells were gated based on FSC-A/FSC-H profiles, and cell-cycle distribution was analyzed using the Watson algorithm in FlowJo.

### Immunofluorescence (IF)

Oocytes, COCs, or ovarian sections were subjected to standard immunofluorescence procedures, including fixation with 4% PFA, permeabilization, blocking, incubation with primary antibodies (Table S1), and fluorophore-conjugated secondary antibodies with DAPI counterstaining. Ovarian sections were additionally subjected to antigen retrieval. Images were acquired using Zeiss LSM880 or LSM710 confocal microscopes, and fluorescence signals were quantified using ImageJ. For F-actin staining, samples were labeled with rhodamine-conjugated phalloidin. Microvilli were visualized by p-ERM immunostaining and Z-stack imaging. Microvilli and transzonal projection (TZP) density were quantified by straightening the oocyte cortex and measuring fluorescence intensity along defined lines using Photoshop and ImageJ. LAMP1 foci were quantified using ImageJ. H3ph10-positive granulosa cells in COCs were manually counted from Z-stack images. For ovarian tissue IF, mean fluorescence intensity was measured within defined regions of interest using ImageJ.

### Live cell staining and imaging of oocytes

For live imaging of mitochondria, ER, and lipid droplets (LDs), FGO were individually stained with Mito-Tracker Red (Thermo, M7512), ER-Tracker Green (APExBIO, B8812), or Nile Red in M2 medium or HBSS containing Ca²⁺ and Mg²⁺ according to the manufacturers’ instructions. Samples were incubated for 30 min at 37°C with Hoechst for nuclear staining. Oocytes were then embedded in 1% low-melting-point agarose containing 5 μM milrinone and imaged using a Zeiss LSM880 confocal microscope at 37°C and 5% CO₂.

### Reconstitution of COCs

COCs were collected from 4-week-old female mice 44–48 h after PMSG injection. Cumulus cell masses were mechanically separated from denuded oocytes using a 90-μm-opening Pasteur pipette. Denuded oocytes and cumulus cell masses were recombined in reconstitution medium containing TCM-199 (80%; Sigma, M4530), sodium pyruvate (0.22%; Sigma, P5280), estrogen (0.8 μg/mL; Sigma, E2758), FSH (4 IU/mL; Merck), FBS (20%; Gibco, 12483020), hyaluronic acid (300 μg/mL; Sigma, 40583), cilostamide (10 μM; Sigma, C7971), and melatonin (10 μM; Sigma, M5250). Reconstituted COCs comprised WT oocytes with *Dhx36oo-/-*cumulus cells, *Dhx36oo-/-*oocytes with WT cumulus cells, or WT oocytes with WT cumulus cells as controls. COCs were cultured for 24, 48, or 60 h, and cumulus expansion was assessed by stereomicroscopy as previously described.

### RNA-seq of cumulus cells

Cumulus cells were collected from female mice 44–48 h after PMSG injection. For each sample, 30–40 COCs were washed with 0.2% BSA in PBS, and cumulus cells were mechanically dissociated using wide-and fine-bore capillaries while minimizing cell loss. Denuded oocytes were promptly removed, and cumulus cells were collected by centrifugation at 3500 × *g* and 4°C and stored at −80°C. SMART-seq2 libraries^50^ were generated by direct lysis, oligo-dT30VN-primed reverse transcription, template switching, and cDNA preamplification. Starting material was normalized to 500 pg cDNA, with amplified cDNA maintained above 5 ng/μL. Libraries were prepared using the TruePrep DNA Library Prep Kit V2 for Illumina (Vazyme, TD503) and sequenced on an Illumina NovaSeq 6000 platform with 150-bp paired-end reads.

Reads were processed using Trim Galore and FastQC, aligned to the mouse mm10 genome using HISAT2, and quantified with featureCounts. Differentially expressed genes were identified using DESeq2 (|log₂FC| ≥ 1, adjusted *P* < 0.05), with FPKM used for visualization and ClusterProfiler for functional enrichment.

### Lipid extraction and quantification by mass spectrometry

Lipids were extracted from 40 FGO using chloroform/methanol (2:1, v/v) as previously described^51,52^ with modifications. Briefly, oocytes were washed with 0.2% non-fat BSA in PBS and quenched in −20°C methanol containing PC 17:0 (1.97 μM), PE 17:0 (2.08 μM), and PS 14:0 (2.14 μM) as spike-in standards. After freeze–thaw lysis, lipids were extracted with chloroform, followed by phase separation with chloroform and 50 mM citric acid. The lipid phase was collected, aliquoted, and dried by vacuum concentration.

Lipid quantification was performed by UHPLC-QTRAP 6500+ mass spectrometry using a multiple reaction monitoring (MRM) method56. Dried extracts were reconstituted in class-specific solvents containing 5 mM ammonium acetate and a deuterated lipid internal standard mixture (Lipidomix, Avanti Polar Lipids, #330707). Lipid species were identified and quantified based on MRM transitions and verified against authentic standards. Peak areas were integrated and manually reviewed using Analyst (SCIEX). Differentially abundant lipids were identified using Limma, followed by lipid metabolic pathway enrichment using a custom R pipeline.

### Promoter G-quadruplex motif analysis

Promoter sequences (2 kb upstream of the transcription start site) of target genes were obtained from the UCSC Genome Browser, and both DNA strands were analyzed using a custom Python script. Potential G-quadruplex-forming sequences (PQS) were identified using the QGRS Mapper^45^ with a minimum sequence length of 30 nt, at least three G-groups, and loop lengths of 1–6 nt. PQS numbers were determined for each promoter and compared across gene sets.

### Evaluation of G-quadruplex formation in vitro by spectra spectroscopy using CYTO

Synthetic PQS oligonucleotides were dissolved in ddH₂O and incubated at 2 μmol/L with 5 μmol/L CYTO in 10 mmol/L Tris-HCl (pH 7.5) containing 60 mmol/L KCl. Fluorescence spectra were recorded from 550–800 nm with excitation at 527 nm using a multi-scan spectrophotometer.

### Western blot

For Western blotting, 100–200 FGO or individual ovaries were lysed in 20 or 100 μL of 2× sample buffer, respectively, with oocyte samples heated at 95°C for 10 min; at least 150 oocytes were used for DHX36 detection. Proteins were separated by SDS-PAGE, transferred to PVDF membranes, blocked with 5% non-fat milk, and incubated with primary antibodies (**Table S1**) overnight at 4°C, followed by HRP-conjugated secondary antibodies. Signals were detected by ECL and captured on X-ray film.

### RT-qPCR

For RT-qPCR analysis of oocytes, 10–20 growing oocytes were collected per sample, washed three times with 0.2% BSA in PBS, and stored at −80°C. Samples were lysed and subjected to random-primer annealing in 4.5 μL of lysis/annealing mixture at 72°C for 3 min, followed by reverse transcription in a 5-μL reaction containing PrimeScript II Reverse Transcriptase (TaKaRa, 2690B) at 30°C for 10 min, 42°C for 60 min, and 75°C for 15 min. cDNA was diluted threefold and analyzed using Power SYBR Green PCR Master Mix (Applied Biosystems, 4367659) on a Bio-Rad CFX96 Touch system. Primer sequences are listed in **Table S2**. *Gapdh* was used as the reference gene, and relative transcript abundance was calculated using the 2^−ΔCt method.

### Transmission electron microscopy (TEM)

Ovaries from 4-week-old mice were dissected into tissue pieces containing intact antral follicles and fixed overnight at 4°C in 2% paraformaldehyde, 2% glutaraldehyde, and 0.1 M sodium cacodylate (pH 7.2; Electron Microscopy Sciences). Samples were post-fixed with 2% OsO_4_ for 2 h, dehydrated, resin-embedded, and polymerized at 65°C for 48 h. Ultrathin sections (90 nm) were prepared, stained with 4% uranyl acetate and 2.5% lead nitrate, and examined by TEM at 80 kV. Microvilli density was quantified as the number of microvilli per 5 μm of oocyte membrane, and TZP density as the number of TZP profiles per randomly selected 3.5-μm² ROI.

### Measurement of serum hormone level

Blood samples were collected from 4-week-old female mice by retro-orbital bleeding under anesthesia at 44 h after PMSG injection or 48 h after hCG administration following PMSG priming. Blood was allowed to clot for 30 min at room temperature and then kept at 4°C overnight. Serum was collected after centrifugation at 12,000 × *g* for 10 min at 4°C. Serum estradiol (E2) and progesterone (P4) levels were measured using the UniCel DxI 800 Access Immunoassay System (Beckman Coulter), with at least 200 μL serum used per measurement.

### Lucifer yellow microinjection into cumulus-oocyte complexes

Dye transfer assay was performed as previously described with modifications^53^. COCs were placed in M2 medium, and cumulus cells on one side of the oocyte were gently removed using a holding pipette. Approximately 7–8 pL of 1 mM Lucifer yellow was microinjected into the oocyte. After 10 min at room temperature, fluorescence images were acquired, and fluorescence intensity was quantified using ImageJ. Dye transfer efficiency was calculated as the ratio of fluorescence intensity in cumulus cells to that in the oocyte.

## Statistical Analysis

All experiments were independently repeated at least three times. Differences between two groups were assessed for statistical significance using a two-tailed Student’s *t*-test. Most of quantitative data are presented as bar plots or violin plots using GraphPad Prism software (version 10). Exact *P* values are annotated directly on the graphs.

