## Supplementary figures and tables for "DHX36 regulates antral follicle development and ovulation as a non-OSF maternal factor by maintaining oocyte homeostasis and supporting OSF delivery"

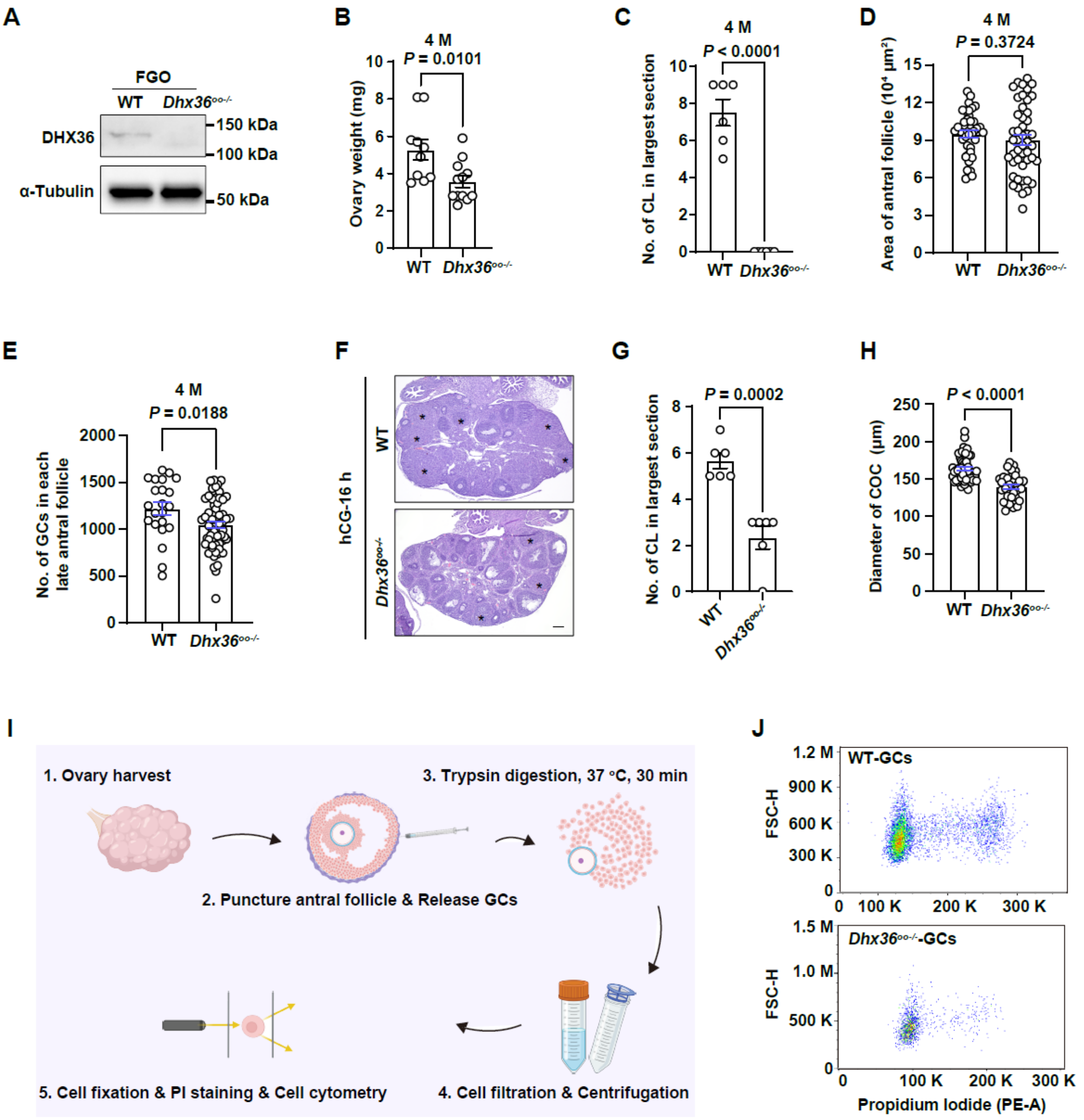


**Figure S1. Oocyte-specific loss of DHX36 impairs granulosa cell proliferation**

**A:** Western blot analysis of DHX36 protein level in FGO isolated from WT and *Dhx36*$^{oo-/-}$ mice. **B:** Ovarian weight of 4-month-old WT and *Dhx36*$^{oo-/-}$ mice. **C:** Number of CL in largest section per ovary in 4-month-old mice. **D:** Quantification of area of antral follicle sections from 4-month-old WT and *Dhx36*$^{oo-/-}$ mice. **E:** Number of granulosa cells in antral follicle section from 4-month-old WT and *Dhx36*$^{oo-/-}$ mice. **F:** H&E staining images of ovaries from 4-week-old WT and *Dhx36*$^{oo-/-}$ mice after hCG (16 h) following treatment with PMSG (44 h). Asterisks indicate CL. (Scale bar: 100 μm). **G:** Number of CL in largest section per ovary in 4-week-old mice following hCG for 16 h. **H:** Diameter quantification of COC from WT and *Dhx36*$^{oo-/-}$ mice. **I:** Schematic diagram of flow cytometry assay for granulosa cells **J:** Flow cytometry analysis of granulosa cells isolated from antral follicles of WT and *Dhx36*$^{oo-/-}$ mice after PMSG treatment for 44 h. Cell populations are displayed on a forward scatter-height (FSC-H**:** vs. propidium iodide intensity plot.

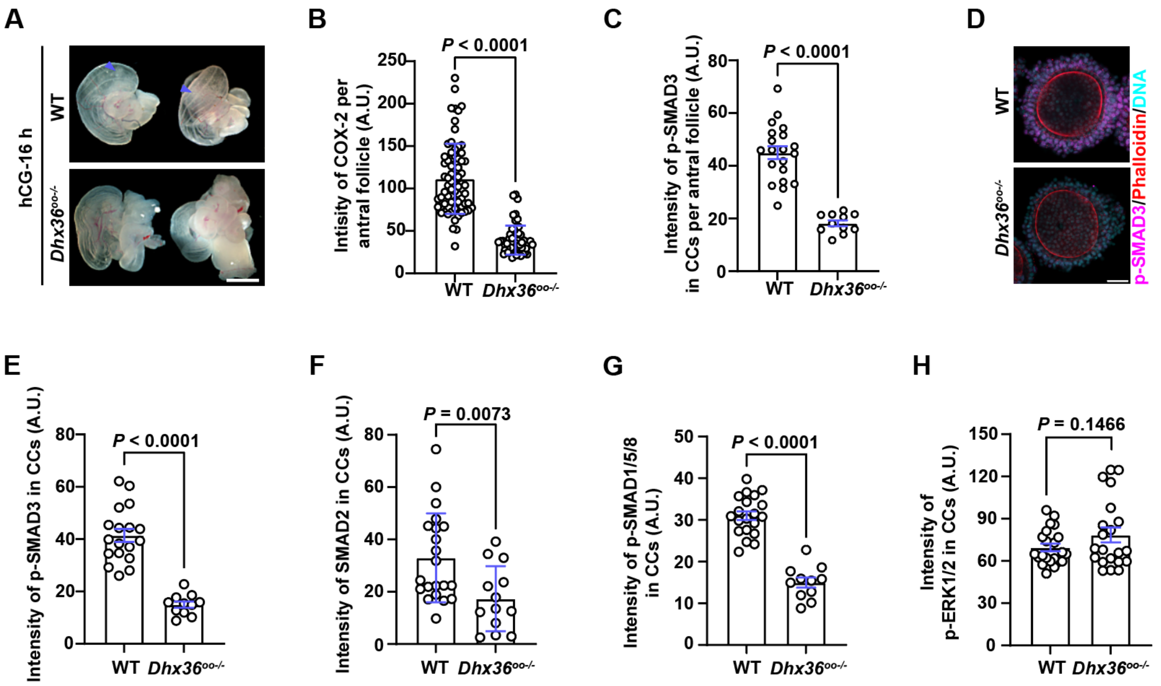


**Figure S2. Impaired SMAD signaling in cumulus cells of *Dhx36*$^{oo-/-}$ mice**

**A:** Representative images of the oviductal ampulla from 4-week-old WT and *Dhx36*$^{oo-/-}$ mice treated with hCG for 16 h following PMSG priming for 44 h. Purple triangles indicate the COCs. (Scale bar: 1 mm). **B:** Quantification of COX-2 fluorescence intensity in antral follicles at 8 h post-hCG (hCG-8 h). **C:** Quantification of p-SMAD3 fluorescence intensity in cumulus cells of antral follicles at 2 h post-hCG (hCG-2 h). **D:** IF staining of p-SMAD3 in COCs collected from 4-week-old WT and *Dhx36*$^{oo-/-}$ mice at hCG-2 h following PMSG priming. (Scale bar: 20 μm). **E–H:** Quantification of fluorescence intensity of p-SMAD3 (E), p-SMAD2 (F), p-SMAD1/5/8 (G), and p-ERK1/2 (H) in cumulus cells of COCs from WT and *Dhx36*$^{oo-/-}$ mice at hCG-2 h. All quantitative data are presented as mean ± SEM. *P* values were calculated using unpaired *t*-tests.

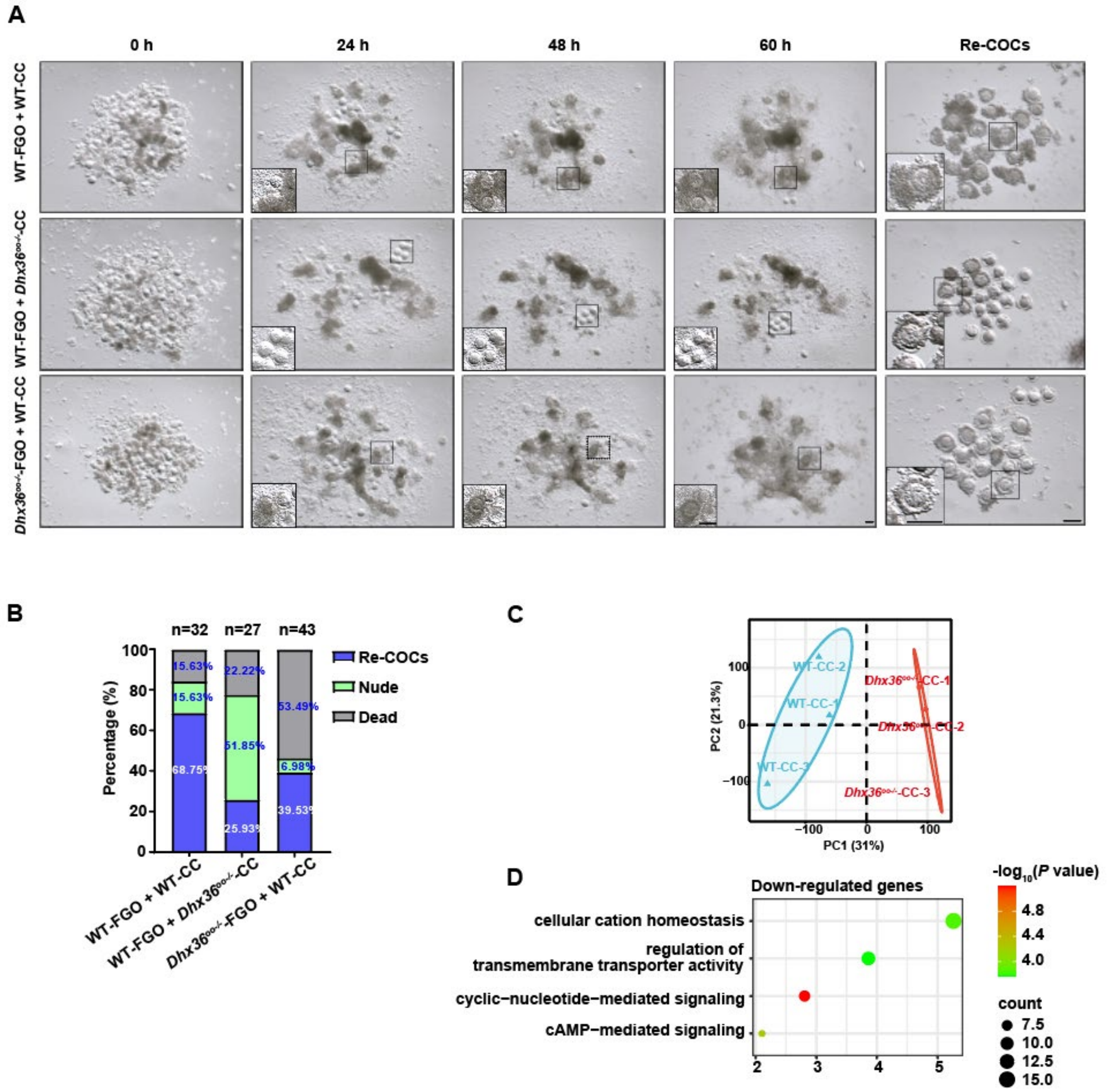


**Figure S3. RNA-seq analysis of cumulus cells from WT and *Dhx36*$^{oo-/-}$ mice**

**A:** Bright-field images of reconstituted COCs from the groups indicated in (C), captured at 0, 24, 48, and 60 h. (Scale bar: 100 μm). **B:** Histogram showing percentage of successfully reconstituted COCs (re-COCs), nude oocytes, and dead oocytes in each group following COC reconstitution. **C:** Principal component analysis (PCA) of RNA-seq data from cumulus cells of 4-week-old WT and *Dhx36*$^{oo-/-}$ mice. **D:** Gene ontology enrichment analysis of biological processes among downregulated genes in cumulus from *Dhx36*$^{oo-/-}$ mice.

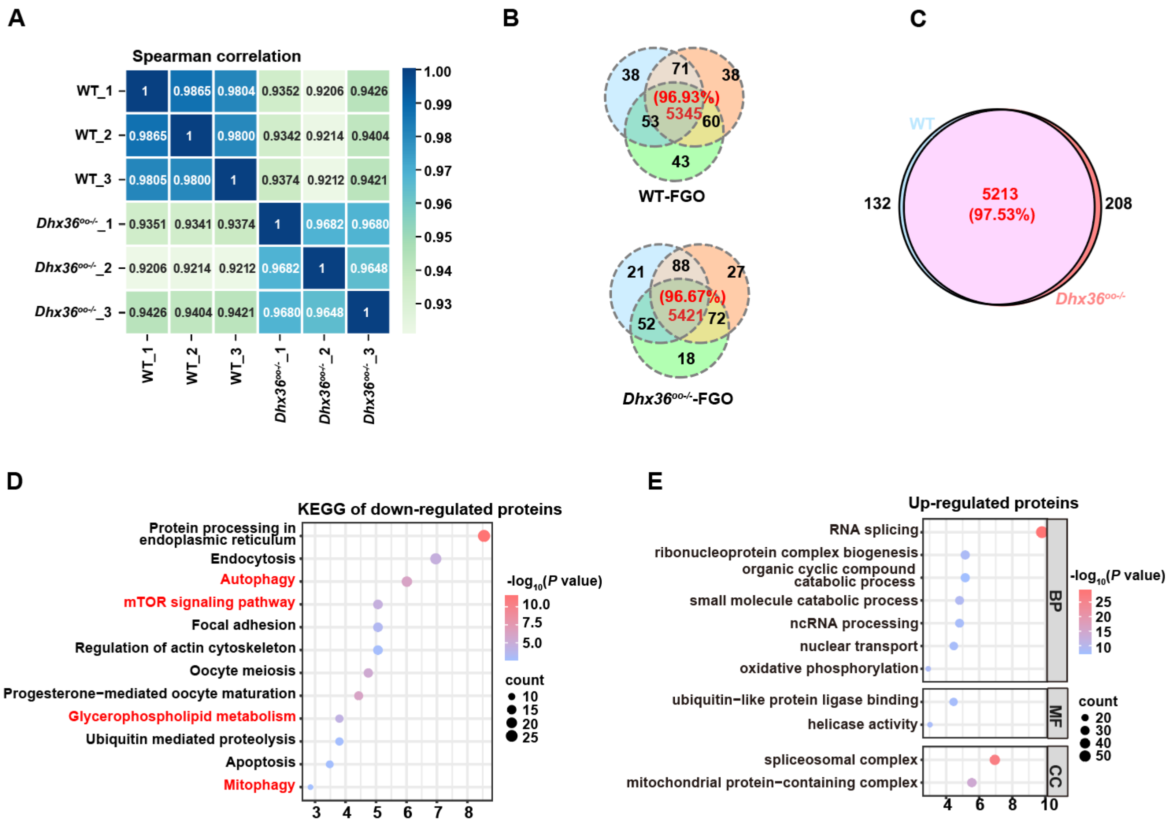


**Figure S4. Additional proteomic data analysis**

**A** Heatmap depicting Spearman correlation between proteomic samples from WT and *Dhx36*$^{oo-/-}$ FGO. **B:** Non-proportional Venn diagram showing the total number of proteins identified in each replicate of WT and *Dhx36*$^{oo-/-}$ FGO. **C:** Proportional Venn diagram showing the overlap of identified proteins between WT and *Dhx36*$^{oo-/-}$ groups. **D:** Kyoto Encyclopedia of Genes and Genomes (KEGG) pathway analysis of downregulated proteins in *Dhx36*-null FGO. **E:** Gene ontology analysis of upregulated proteins in *Dhx36*-null FGO.

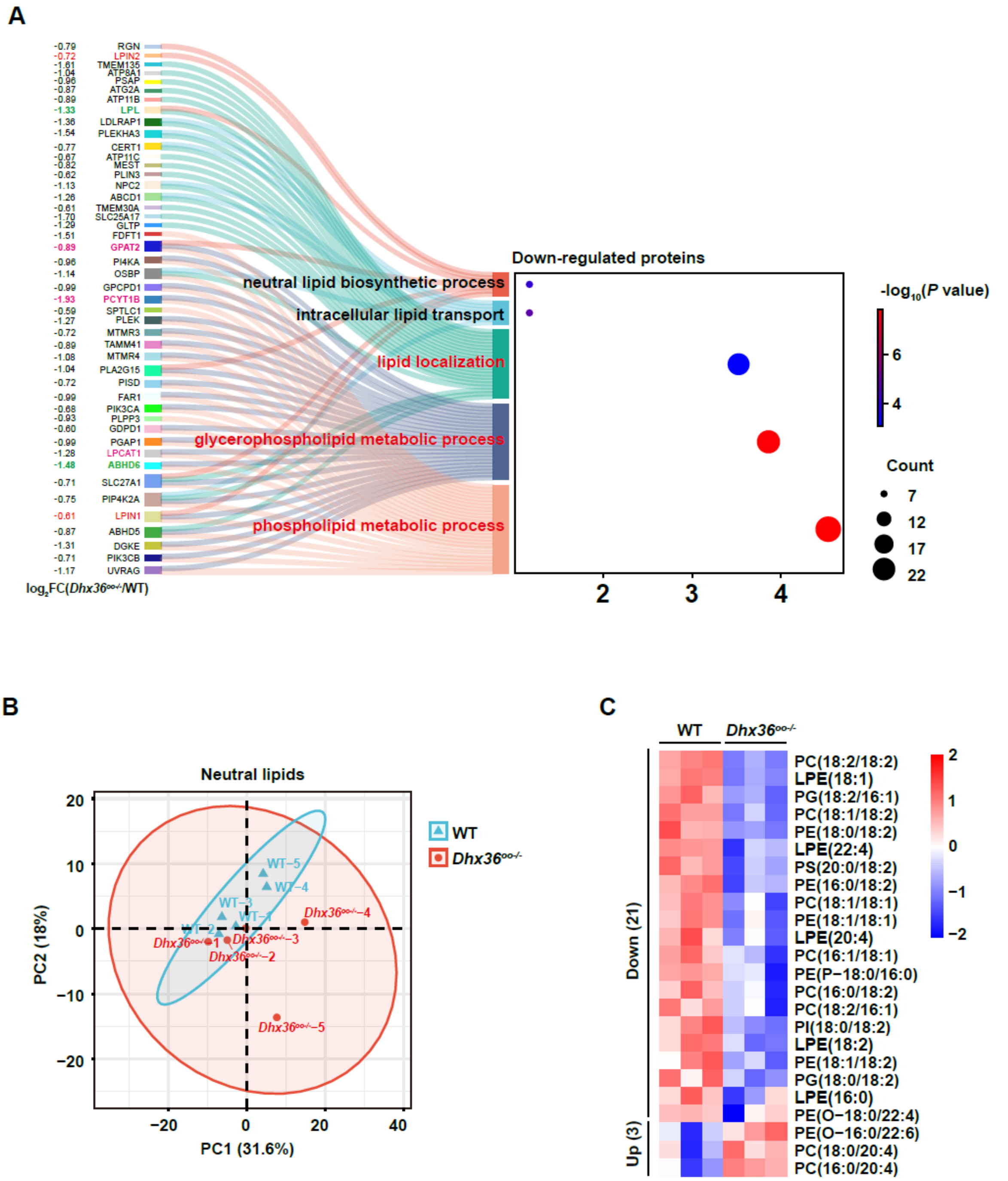


**Figure S5. Additional analysis of lipidomic data**

**A:** Integrated bubble and Sankey plot showing downregulated proteins in *Dhx36*$^{oo-/-}$ FGO enriched in lipid metabolism pathways. **B:** PCA of neutral lipid profiles in FGO from WT and *Dhx36*$^{oo-/-}$ mice. **C:** Heatmap of differentially abundant phospholipids in *Dhx36*$^{oo-/-}$ versus WT FGO. Significance criteria were designated as $|\log_2 FC| \geq 0.263$ (equivalent to ≥ 1.2-fold change) and *P* value < 0.05 ($-\log_{10}(P) \geq 1.30$).

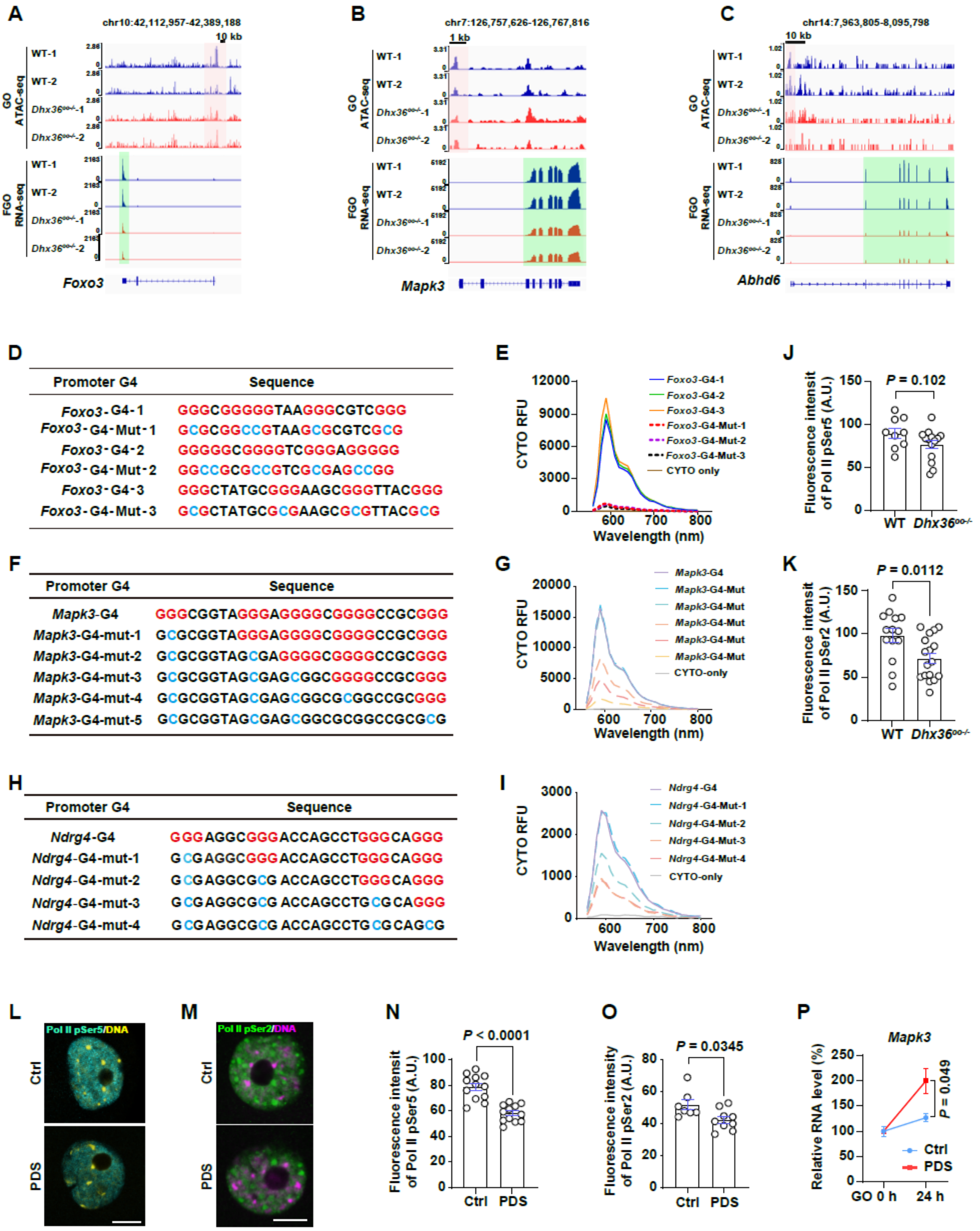


**Figure S6. G4 stabilization suppresses polymerase II activity in oocytes**

**A–C:** IGV tracks showing ATAC-seq and RNA-seq data for *Foxo3* (A) and *Mapk3* (B) in growing oocytes (GO) and FGO from WT and *Dhx36*$^{oo-/-}$ mice. **D, F, H:** Tables listing wild-type and mutant PQS within the promoters of *Foxo3* (D), *Mapk3* (F) and *Ndrg4* (H). **E, G, I:**

Fluorescence spectra of the PQS and their corresponding mutants listed in (D, F, H) after incubation with G4 fluorescent probe CYTO. **J, K:** Quantification of fluorescence intensity of Pol II pSer5 (J) and Pol II pSer2 (K) in growing oocytes from WT and *Dhx36*$^{oo-/-}$ mice. **L, M:** Immunofluorescence staining of Pol II pSer5 (L) and Pol II pSer2 (M) in growing oocytes treated with or without 10 μM PDS for 24 h. (Scale bar: 20 μm). **N, O:** Quantification of fluorescence intensity of Pol II pSer5 (N) and Pol II pSer2 (O) in growing oocytes with or without PDS treatment for 24 h. **P:** RNA level of *Mapk3* in growing oocytes before and after PDS treatment (10 μM) for 24 h, as detected by RT-qPCR. All quantitative data are presented as mean ± SEM. *P* values were calculated by unpaired *t*-tests.

**Table S1. The information of primary antibodies.**

| Antibody | Manufacture | Catalogue | Dilution |
|---|---|---|---|
| anti-DDB1 (Rabbit) | Abcam | ab109027 | WB (1:1000) |
| anti-DHX36 (Rabbit) | Abcam | ab70269 | WB (1:1500) |
| p44/42 MAPK (ERK1/2) (Rabbit) | CST | 4695 | WB (1:1000) |
| anti-p-ERK1/2 (Rabbit) | CST | 9101 | WB (1:1000) |
| anti-p-ERM (Rabbit) | CST | mAb#3726 | IF (1:100) |
| anti-p-SMAD1/5/8 (Rabbit) | CST | 9511S | WB (1:1000)<br>IF (1:100) |
| anti-p-SMAD2/3 (Rabbit) | CST | 9510 | WB (1:1000) |
| anti-p-SMAD2 (Rabbit) | | | IF (1:100) |
| anti-p-SMAD3 (Rabbit) | Selleckchem | F2846 | IF (1:100) |
| anti-GDF-9 (Rabbit) | Proteintech | 29309-1-AP | WB (1:1000)<br>IF (1:200) |
| anti-BMP-15 (Rabbit) | Proteintech | 18981-1-AP | WB (1:1000)<br>IF (1:200) |
| anti-H3ph10 (Rabbit) | CST | 9701S | IHC (1:200)<br>IF (1:200) |

| Antibody | Manufacture | Catalogue | Dilution |
|---|---|---|---|
| anti-BrdU (Mouse) | Sigma-Aldrich | B2531 | IHC (1:1000) |
| anti-LAMP1 (Rabbit) | CST | E5N9Z | IF (1:200) |
| anti-COX-2 (Rabbit) | Santa Cruz | sc-19999 | WB (1:1000)<br>IF (1:200) |
| anti-PTX3 (Goat) | R&D System | AF2166 | WB (1:1000)<br>IF (1:200) |
| Anti-RNA polymerase II CTD repeat YSPTSPS (phospho S2) (PS2) | Abcam | ab193468) | IF (1:200) |
| anti-RNA polymerase II CTD repeat YSPTSPS (phospho S5) (PS5) | Abcam | ab5131 | IF (1:200) |

**Table S2. The RT-qPCR primers.**

| Name | Sequence (5’ to 3’) |
|---|---|
| *Gapdh*-F | ACACTGAGGACCAGGTTGTCTC |
| *Gapdh*-R | TACTCCTTGGAGGCCATGTAG |
| *Foxo3*-F | AAGGGAAGGAGCCGAGGTAG |
| *Foxo3*-R | CTCTGTGGCTCGAACTCTGG |
| *Ndrg4*-F | CCATCTCTTCAGCCAGGAGG |
| *Ndrg4*-R | TCCAGAAGAGCTGCAGGTTG |
| *Mapk3*-F | GTACGGCATGGTCAGCTCAG |
| *Mapk3*-R | CTGAGGATGTCTCGGATGCC |

**Table S3. Summary of genes used for G4 motif analysis**

| type | gene number | total number with G4 in both strands | number of genes with G4 | percentage of genes with G4(%) |
|---|---|---|---|---|
| All genes | 5031 | 3066 | 2039 | 40.52 |
| Organelle organization genes | 26 | 18 | 12 | 46.15 |
| Lipid metabolism genes | 14 | 14 | 9 | 64.28 |
| Membrane assembly genes | 12 | 10 | 7 | 58.33 |
| Autophagy genes | 6 | 5 | 3 | 50.00 |

**Table S4 RNA-seq data of cumulus cells from WT and *Dhx36*$^{oo-/-}$ mice.**

**Table S5 Proteomics data of FGO from WT and *Dhx36*$^{oo-/-}$ mice.**

**Table S6 Lipid metabolism data of FGO from WT and *Dhx36*$^{oo-/-}$ mice.**
